# Peroxiredoxin 2 mediates redox-stimulated adaptations to oxidative phosphorylation induced by contractile activity in human skeletal muscle myotubes

**DOI:** 10.1101/2024.09.23.611634

**Authors:** Robert A Heaton, Sam T M Ball, Caroline A Staunton, Vincent Mouly, Samantha W Jones, Anne McArdle, Malcolm J Jackson

**Affiliations:** MRC-Versus Arthritis Centre for Integrated Research into Musculoskeletal Ageing (CIMA), Department of Musculoskeletal and Ageing Science, Institute of Life Course and Medical Sciences, University of Liverpool, Liverpool L7 8TX, U.K; URA CNRS 1448, UFR Biomédicale des St. Pères, Paris, France

## Abstract

Skeletal muscle generates superoxide during contractions, which is converted to hydrogen peroxide (H_2_O_2_). H_2_O_2_ has been proposed to activate signalling pathways and transcription factors that regulate adaptive responses to exercise, but the concentration required to oxidize and activate key redox-sensitive signalling proteins *in vitro* is much higher than the typical intracellular levels seen in muscle after exercise. We hypothesized that 2-Cys-peroxiredoxins (PRDX), which rapidly oxidize in the presence of physiological concentrations of H_2_O_2_, serve as intermediary signalling molecules and play a crucial role in activating adaptive pathways following muscle contractions. This study has examined the human muscle myotube responses to contractile activity, or exposure to low extracellular concentrations (2.5-5 µM) of H_2_O_2_ and whether knock down of muscle PRDX2 alters the differential gene expression (DEG) that results from these stresses. Exposure of human skeletal muscle myotubes to a 15 min period of aerobic electrically stimulated isometric contractions or 5μM H_2_O_2_ induced substantial changes in DEG with modification of many genes associated with adaptations of skeletal muscle to contractile activity. Common DEG in these conditions included upregulation of genes associated with increased mitochondrial oxidative phosphorylation, including *COX1, COX2, COX3* and *ATP6*. In myotubes with PRDX2 knock down (94% decrease in PRDX2 mRNA), the upregulation of genes associated with increased mitochondrial oxidative phosphorylation was abolished following contractile activity or exposure to H_2_O_2_. These data indicate that a common effect of contractile activity and exposure to “physiological” levels of H_2_O_2_ in human myotubes is to increase the expression of multiple genes associated with increased mitochondrial oxidative phosphorylation. Furthermore, these effects were abolished in PRDX2 knock down myotubes indicating that adaptations to upregulate multiple genes related to increased mitochondrial capacity in human muscle myotubes in response to exercise is both redox regulated and requires PRDX2 as an essential mediator of the effects of H_2_O_2_.

## Introduction

The contractile activity of skeletal muscle results in the generation of superoxide and nitric oxide leading to the formation of secondary reactive oxygen species (ROS), such as hydrogen peroxide (H_2_O_2_), and reactive nitrogen species [1–4]. Increased H_2_O_2_ generation has been linked to stimulation of adaptive responses of muscle to repeat contractions via activation of a number of signalling pathways and transcription factors leading to the upregulation of genes involved in stress responses, catabolism, glucose uptake and mitochondrial biogenesis [5–11]. The extent of the adaptations mediated by increased H_2_O_2_ production is unclear, although some previous studies examining the effect of high dose antioxidants in exercising human or animal subjects suggest that the pathways stimulated by oxidants may be extensive [12]. One issue which has complicated the identification of the exact processes involved is the discrepancy between the very low physiological *in vivo* concentrations of intracellular H_2_O_2_ and those reported to activate known redox activated signalling proteins and transcription factors *in vitro* [13]. We have previously calculated that the cytosolic concentration of H_2_O_2_ increases to around 10 M in muscle fibres following contractile activity [14, 15] although these figures may be an overestimate due to revised analysis of cellular H_2_O_2_ kinetics [16]. The low intracellular H_2_O_2_ concentrations found in the cytosol are thought to be insufficient to directly oxidise known redox-sensitive proteins in key signalling pathways or transcription factors.

To explain this discrepancy in how changes in the low cytosolic levels of H_2_O_2_ can activate less sensitive redox-signalling proteins the process of redox-relays has been proposed in which H_2_O_2_ initially oxidises thiol peroxidases, such as the 2-Cys peroxiredoxins (PRDX), as mediators of subsequent oxidation of other protein thiols [17]. PRDX are present in multiple cellular compartments and are able to scavenge and react with H_2_O_2_ in cells at very low physiological concentrations and are several orders of magnitude more reactive with H_2_O_2_ than many other proteins reported to be redox-regulated [18, 19]. PRDX play an intermediary role in transferring H_2_O_2_-derived oxidising equivalents to a number of redox regulated target proteins [20–24]. The precise mechanism by which this occurs is unclear, but some of these target proteins are capable of forming heterodimers with PRDX becoming oxidised and initiating the “redox relay” [24], PRDX’s are therefore proteins that potentially link contraction-induced H_2_O_2_ generation with activation of multiple redox-regulated signalling pathways.

In previous studies we have shown that a short exposure of isolated mature murine skeletal muscle fibres to extracellular concentrations of H_2_O_2_ as low as 2.5µM, or a short period of isometric contractile activity, leads to oxidation and formation of homodimers of PRDX1, 2 and 3. The contraction-induced oxidation of PRDX2 occurring rapidly within 1 minute (after 12 maximal contractions) [26]. Such data suggests that PRDX1, 2 or 3 may be effectors of skeletal muscle redox signalling following contractions. Subsequently Xia et al have demonstrated that in *C. elegans* deletion of PRDX2 caused decreased adaptations following swimming activity and that PRDX2 potentially acts via DAF-16 to mediate mitochondrial remodelling [27, 28].

In the current studies we have examined the effect of deletion of PRDX2 in human skeletal muscle myotubes. PRDX2 was selected for study since it was rapidly oxidised during contractile activity in our previous studies [26] and the muscle content of PRDX2 exceeds that of PRDXs1 or 3[29]. Control myotubes (transduced with a scrambled shRNA) and myotubes in which PRDX2 was substantially deleted (PRDXKD myotubes) were treated with “physiological” levels of H_2_O_2_ (2.5 or 5µM) or electrically stimulated to contract and the gene expression changes analysed using an unbiased approach with RNA deep sequencing (RNAseq) analyses. We hypothesised that examination of the differential gene expression (DEG) between experimental groups would allow identification of those genes and adaptive pathways that were altered by both contractile activity and H_2_O_2_ and hence were potentially redox-regulated. Furthermore, comparisons with the DEG in PRDX2 Knockdown (PRDX2KD) myotubes following exposure to either contractile activity or H_2_O_2_ would allow us to identify those adaptive pathways likely to be mediated by PRDX2.

## Materials and Methods

### Cell Culture

The human skeletal muscle cell line AB1167 derived from the *Fascia lata* of a 20-year-old male was used in this study and was provided by Genethon (https://www.genethon.com/) [30]. Myotubes derived from this line demonstrate a phenotype for contractions both spontaneously and following electrical stimulation making them suitable muscle cell for the current study. Myoblasts were grown using a medium consisting of 20% Medium 199, GlutaMAX (Invitrogen, Thermo Fisher Scientific, Waltham, MA, USA) and 80% DMEM, GlutaMAX (Gibco, Thermo Fisher Scientific, Waltham, MA, USA). This medium was supplemented with 10% Foetal Bovine Serum (ThermoFisher, Waltham, MA, USA), 0.4 µg/ml Dexamethasone (Sigma-Aldrich, St. Louis, MO, USA), 5 ng/ml hEGF (Gibco, Thermo Fisher Scientific, Waltham, MA, USA), 0.5 ng/ml bFGF (Gibco, Thermo Fisher Scientific, Waltham, MA, USA ), 25 µg/ml Fetuin (Sigma-Aldrich, St. Louis, MO, USA), 50 µg/ml Gentamicin (Sigma-Aldrich, St. Louis, MO, USA), and 5 µg/ml Insulin (Sigma-Aldrich, St. Louis, MO, USA) and kept at 37°C and 6% CO_2_. Differentiation of myoblasts to myotubes was conducted using DMEM (Invitrogen, 61965-026), containing 25 mM glucose with GlutaMAX formulation (lacking sodium pyruvate, Gibco, Thermo Fisher Scientific, Waltham, MA, USA). The media was supplemented with 1% of gentamicin (50 µg/ml), insulin (10 µg/ml), and x 2% (10ml) of horse serum. Cells were approximately 80% confluent prior to adding differentiation medium (DM). After adding DM, cells were kept at 37°C and 6% CO_2_ for a minimum of five days to induce differentiation into myotubes which were used for experimental procedures. Methods for tissue culture and differentiation were adapted from Furling [31].

For western blot analysis, myotubes were washed using PBS then harvested by adding 50 μl Radioimmunoprecipitation Assay (RIPA) buffer containing protease inhibitors (Roche, Basel, Switzerland). Myotubes where scraped from the well and lysates centrifuged at 12000g for 5 min and supernatant collected. The total protein concentration was determined using the Bradford assay [32].

Harvesting for both rt-QPCR and RNA sequencing was undertaken as follows. DM media was removed, and myotubes washed twice with PBS. The cells were harvested via scraping into the D-PBS and cells were transferred to an Eppendorf tube and centrifuged as above and supernatant removed. The pellet was then re-suspended in 350 μl of Qiagen RLT lysis buffer and disrupted by gentle pipetting. Three hundred and fifty μl of 70% ethanol was added and RNA was extracted using the Qiagen mini-prep RNA extraction kit following manufacturer’s instructions. Extracted RNA was stored in 30µl of RNA free purified water at -80°C until needed.

### Knock down of PRDX2 in myoblasts

To achieve knockdown of PRDX2 in myoblasts, a specific shRNA was designed and procured (VectorBuilder, Guangzhou, China). A recombinant adeno-associated virus (rAAV6) was utilized for the delivery of the shRNA into the myoblasts [33] and a mCherry tag was integrated into the vector to serve as a reporter for transduction efficiency. Control myoblasts were transduced with a scrambled PRDX2 shRNA.

For the transduction process, myoblasts were seeded at a density of 250,000 cells per well (6 well plate), in DM without gentamicin. Following plating, the myoblasts were incubated for 24 hours (37°C, 5% CO_2_). Cell adherence to the well was visually confirmed, and the culture medium was carefully aspirated. The myoblasts were then washed twice with PBS to remove any residual medium. Following the wash steps, 1.5 ml of fresh DM containing the rAAV6-shRNA was added to each well. Cells were then incubated in this media for 24 hours, before topping up to 2 ml with fresh DM without gentamicin. At 48 hours post transduction media was removed and replaced with fresh DM media, cells were be maintained in DM for 10 days prior to experimental procedures to allow full myotube differentiation and PRDX2 protein turnover.

The volume of rAAV6 to be added was calculated using a formula considering the desired viral titre, concentration, and the number of cells plated. This was calculated to be between 10 and 10 viral genomes (vg) pec cell in the plate, this calculation ensures an optimal transduction efficiency while not eliciting an immune response based on the work of Chamberlain and colleagues [34]and additional studies undertaken in our laboratory [35].

### Treatments of myotubes

For H_2_O_2_ treatments, media was removed from myotubes and replaced with fresh serum-free medium containing the desired final concentration of 2.5uM or 5uM H_2_O_2_ for 15 minutes as previously [26]. Cells were then washed using fresh PBS which was removed and the myotubes were washed again with D-PBS before harvesting and freezing for storage.

For electrical stimulation experiments, myotubes were placed in 1 ml fresh serum-free medium and electrically stimulated to contract using field stimulation following the protocol described by Jackson and colleagues [2]. Briefly, platinum electrodes were immersed in the culture medium and delivered trains of biphasic square wave pulses lasting 2ms, with a duration of 0.5s, repeated every 5s at 50Hz and 30V for 15 minutes. Myotubes were then harvested immediately following stimulation in an identical procedure to that used following H_2_O_2_ exposure.

### Western blotting analyses

Thirty micrograms of protein from the cell lysate in RIPA buffer was separated by reducing SDS-PAGE on a 4-15% polyacrylamide gel (Bio-Rad, Hercules, California, USA). The protein was transferred to a nitrocellulose membrane using a semi-dry transfer method [36]. The membrane was blocked in 5% milk powder diluted in tris-buffered saline (TBS) for one hour at room temperature. Following blocking, the membranes were incubated overnight at 4°C with the required primary antibody, diluted 1:1000, in 5% powered milk diluted in TBS with 0.1% Tween-20 (TBST) on a rocking table at 4°C.

The membrane was washed 5 x 2 minutes with TBST and incubated with IRDye® 800CW goat anti-rabbit IgG (H + L) secondary antibody (Licor Biosciences, Lincoln, Nebraska, USA), diluted 1:10,000 in 5% milk powder TBST for one hour at room temperature. Subsequently, the membrane was washed 5 x 2 minutes with TBST, before a finally being replaced with H_2_O for imaging. Membranes were imaged using the Odyssey CLx Imaging System (Licor Biosciences, Lincoln, Nebraska, USA), and densitometric analysis was undertaken using Image J. The antibodies used were PRDX2 (Abcam, Cambridge, United Kingdom: ab109367) and GAPDH (Sigma, St. Louis, Missouri, USA: G9545).

### RT-qPCR analysis of PRDX2 mRNA

RNA extraction was undertaken as outlined above, cDNA synthesis was carried out using the BIORAD iScript cDNA synthesis kit, following the manufacturer’s instructions. Amplification of housekeeping and genes of interest was conducted using the ROACH LC96 light cycler, employing SYBR Green for detection. Primer sequences used: PRDX2: forward -CCTTCCAGTACACAGACGAGCA, reverse - CTCACTATCCGTTAGCCAGCCT; β-actin: forward - TTCCAATATGAGATGCGTTGTT, reverse – GCTATCACCTCCCCTGTGTG.

### RNA isolation and library preparation for sequencing

For RNA sequencing, samples were extracted as described above and total RNA was depleted from samples using Ribo-Zero rRNA removal kit, following the manufacturer’s instructions. DNA free RNA was used for polyadenylation using the NEB Next poly(A) mRNA magnetic isolation module (New England Biolabs, Ipswich, UK). Following enrichment, RNA-Seq libraries were prepared using the NEBNext® UltraTM directional RNA library prep kit for Illumina ®#E7420. Subsequent purification was performed using AMPure XP beads (Beckman Coulter, California, USA). Quantification of all libraries was conducted using a Qubit (Thermo Fisher Scientific, Waltham, MA, USA), while size distribution analysis was carried out using the Agilent 2100 Bioanalyzer (Agilent, California, USA). For the DNA samples, quantification was undertaken using the Qubit® dsDNA HS Assay Kit, accompanied by an assessment of quality and average size using the High Sensitivity DNA Kit. Further, a qPCR assay, specifically targeting adapter sequences flanking the Illumina libraries, was performed using the Illumina® KAPA Library Quantification Kit (Kapa Biosystems, Wilmington, USA). The RNA libraries underwent sequencing on an Illumina® HiSeq 4000 platform with version 1 chemistry, employing sequencing by synthesis (SBS) technology to generate 2⍰×⍰150⍰bp paired-end reads. This sequencing procedure was carried out at the Centre for Genomic Research, University of Liverpool (https://www.liverpool.ac.uk/genomic-research/).

### Bioinformatic analysis and read alignment

Base-scaling and de-multiplexing of indexed reads were executed through CASAVA version 1.8.2 (Illumina) to generate FASTQ format files. Subsequently, the raw FASTQ files underwent trimming to eliminate Illumina adapter sequences, employing Cutadapt version 1.2.1 [37]. Further refinement involved the removal of low-quality bases using Sickle version 1.200, setting a minimum window quality score of 20. Post-trimming reads shorter than 20 bp were excluded. The alignment of reads to the genome sequences was accomplished using TopHat version 2.1.0 [38], followed by transcript assembly conducted through HTSeq2.

### Differential gene expression

Raw gene counts were subject to statistical analysis using an in-house Python pipeline (https://github.com/stmball/prdx/tree/main/prdx). This follows the industry standard DESeq2 package set out by Huber and colleagues [39] to identify the differentially expressed genes (DEGs). Pairwise comparisons were used to identify the DEGs across the treatment groups. The Matplotlib library was used to generate appropriate figures following statistical analysis including volcano plots and Venn diagrams. Fold change was used to identify functional changes considering the false discovery rate where appropriate (FDR)-adjusted p values < 0.05; both Raw and adjusted p-values are reported. All pathway analysis was performed using adjusted p-values. This analysis was conducted for enrichment studies, for Gene Ontology (GO) Biological Process and the “enrichKEGG” package from clusterProfiler v. 3.8.1 in Bioconductor v. 3.7 for Kyoto Encyclopedia of Genes and Genomes (KEGG) pathway enrichment analysis and MSigDB gene set enrichment analysis (Hallmark), with Human as the reference organism. Significantly enriched GO terms and KEGG pathways were identified based on FDR < 0.05.

To determine interaction alterations between groups, pathways, up/downstream regulators and network analysis was performed on the calculated DEGs. This was undertaken using the Ingenuity Pathway Analysis package (IPA) “core analysis” setting (IPA, Qiagen Redwood City, USA). The fundamental analysis was conducted using settings that accounted for both indirect and direct relationships between molecules, relying on experimentally observed data. The Human databases within the Ingenuity Knowledge Base were consulted for data sources. The network scores, determined by the Ingenuity Pathway Analysis (IPA), were computed as the negative exponent of the p-value associated with each network, following the methodology outlined by Tschoeke and colleagues[40].

## Results

### Confirmation of knock down of PRDX2 in human skeletal muscle myotubes

Deletion of PRDX2 in the shRNA treated myotubes was visualised using the mCherry tag, and then confirmed using both western blot and rt-qPCR. A representative western blot confirming the successful knockdown of PRDX2 protein in the human skeletal muscle myotubes at 10 days post transduction compared with scramble-treated (“Scram”) control is shown in Figure 1A. Figure 1B shows PRDX2 mRNA contents comparing Scram control myotubes, and PRDX2KD myotubes. There was a decrease of 94% in PRDX2KD myotubes compared with the Scram control myotubes indicating successful knockdown.

**Figure 1.**
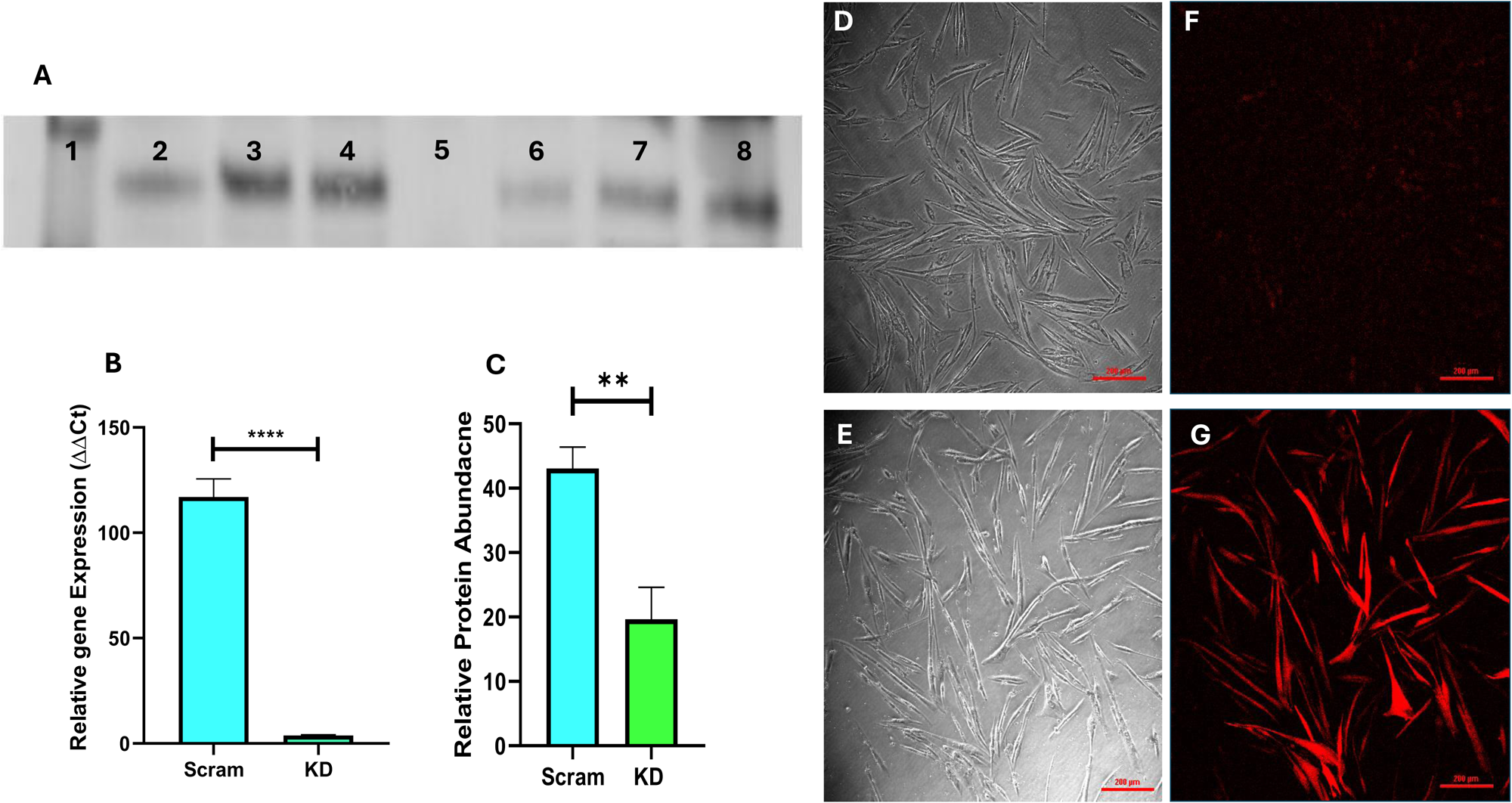
Confirmation of PRDX2 knockdown in human skeletal muscle myotubes. (A) Western blot analysis, the protein contents were quantified using densitometry on Image-J. Key: Lane 1 – Molecular weight marker, Lanes 2-4 - Scram control myotubes, Lane 5 – empty, Lane 6-8 - PRDX2KD myotubes. rt-QPCR relative gene expression levels to scramble control quantified using the delta-delta CT method comparing Scram and KD myotubes (****p<0.0001). (C) Relative protein abundance, comparing cells treated with scrambled shRNA (Scram) and shRNA for PRDX2 (KD) (**p=0.008). Panels D & F show microscopy images of cells post transfection, Panels E & G show fluorescent confocal images post-transfection. D and E show myotubes transfected with the scrambled shRNA, with E showing no increase in M-cherry fluorescence. F and G show myotubes transfected with PRDX2KD shRNA, with G showing myotubes with increased red M-Cherry florescence indicating successful transfection.

The PRDX2 protein quantification comparing the relative protein abundance in the myotubes treated with the scrambled sequence and shRNA (PRDX2KD) treated is shown in Figure 1C. These data show an average 45% decrease in PRDX2 protein content in the PRDX2KD samples compared with the Scram samples.

Figure 1 E and G show representative confocal images of the muscle myotubes 10 days post-transduction with the scramble shRNA. Figure 1E shows little to no fluorescence from the mCherry tag indicating that treatment with the scrambled sequence did not promote the copying of the shRNA plasmid. Figures 1 G illustrate successful transduction of the shRNA PRDX2 plasmid into the PRDX2KD myotubes indicated by the red fluorescence from the mCherry tag (Figure 1G).

Western blotting analysis of the PRDX1 and 3 contents of transduced myotubes showed no significant differences in the protein contents between Scram control and PRDXKD myotubes (data not shown in detail).

### Effect of knock down of PRDX2 on differentially expressed genes (DEGs) in human muscle myotubes at rest

A comprehensive analysis of RNA sequence data was performed to compare Scram control muscle myotubes with PRDX2KD myotubes at rest. The volcano plot presented in Figure 2A shows the extensive differentially expressed genes (DEG) between the 2 groups. The genes that were significantly downregulated in PRDX2KD myotubes compared with scramble control myotubes (548 genes) are indicated in blue, while the significantly upregulated genes are shown in red (541 genes, from 21990 genes in total). Figure 2B shows bar diagrams illustrating the relative changes in expression of some key genes modified in the PRDX2KD myotubes compared with Scram control myotubes. These plots show multiple significant upregulation (red bars) and down regulation (blue bars) of gene expression in the PRDX2KD myotubes, with majority of mitochondrial associated genes being down regulated following PRDX2KD.

**Figure 2.**
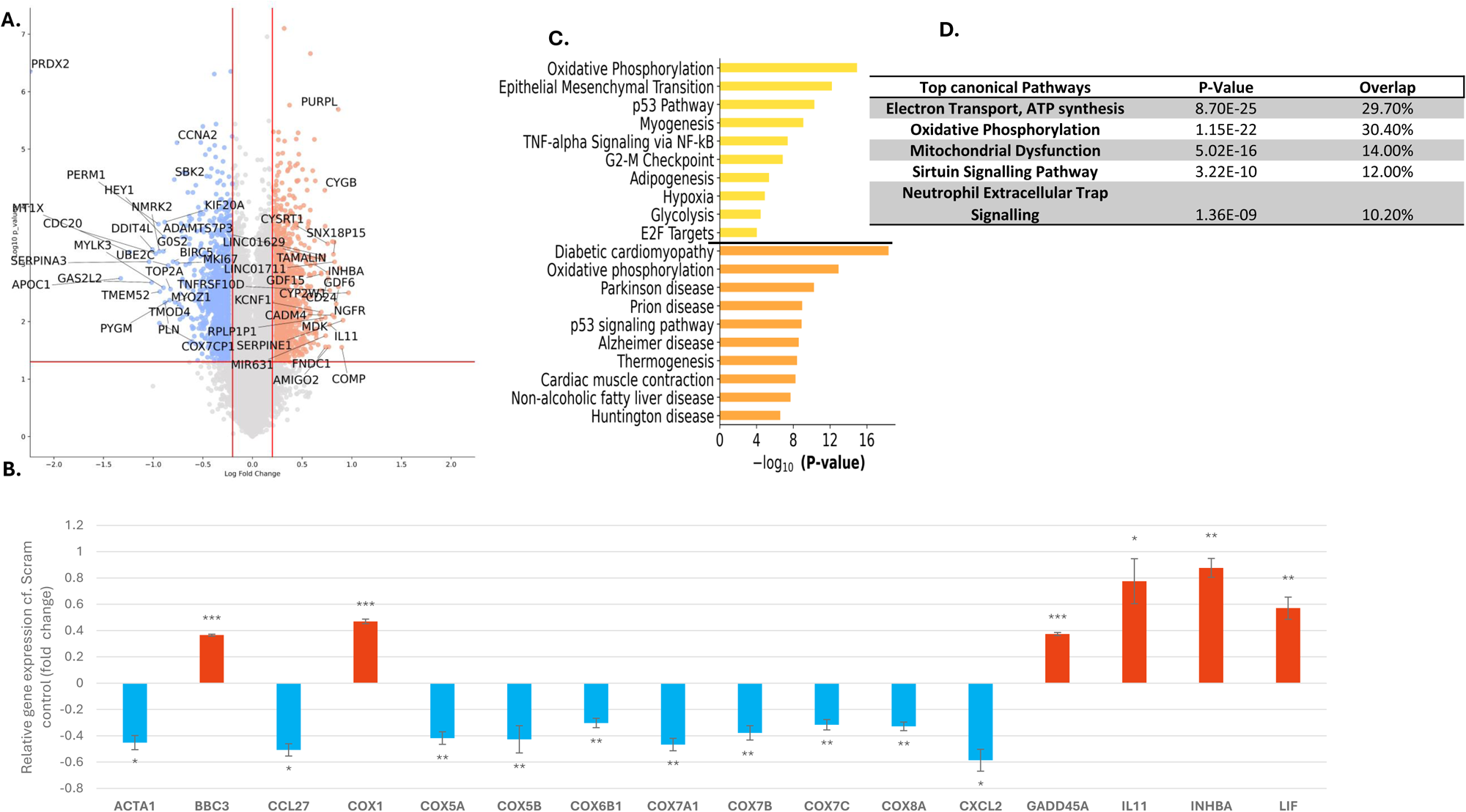
Effect of knock down of PRDX2 on DEGs in human muscle myotubes at rest compared with scram control at rest. (A) Volcano plot showing the differential expression of genes between Scram control and PRDX2KD myotubes. The x-axis indicates the log2 fold change, while the y-axis represents the -log10 (p-value). Genes with significant differential expression are highlighted in blue (downregulated) and red (upregulated). (B) Bar graphs showing the relative changes in expression of key genes discussed in the text. Upregulated genes shown in red with downregulated shown in blue. (C) KEGG (orange) and Hallmark (yellow) pathway analyses highlighting the pathways linked to the gene expression changes (D) IPA analysis of the top five canonical pathways. Gene Key: ACTA1 – Actin, alpha 1, skeletal muscle; BBC3 - BCL2 binding component 3 (also known as PUMA); CCL27 - C-C motif chemokine ligand 27; COX1 - Cytochrome c oxidase subunit 1; COX5A - Cytochrome c oxidase subunit 5A; COX5B - Cytochrome c oxidase subunit 5B; COX6B1 - Cytochrome c oxidase subunit 6B1; COX7A1 Cytochrome c oxidase subunit 7A1 (muscle); COX7C - Cytochrome c oxidase subunit 7C; COX8A - Cytochrome c oxidase subunit 8A (heart/muscle); CXCL2 - C-X-C motif chemokine ligand 2; GADD45A Growth arrest and DNA damage inducible alpha; IL11 – Interleukin 11; INHBA – Inhibin subunit beta A; LIF – Leukaemia inhibitory factor.

The top enriched pathways identified as affected by PRDX2KD using KEGG (yellow) and Hallmark (orange) analyses are shown in Figure 2C. Both analyses show a major effect of the PRDX2KD on genes involved in oxidative phosphorylation. IPA analysis also indicates the relationships between the key genes, pathways and functions that have been altered in PRDX2KD myotubes (Figure 2D). Overall, these analyses indicate that the PRDX2KD influences multiple DEGs linked to mitochondrial function, sirtuin signalling and inflammatory signalling. The IPA analysis also identified TGFβ1 and TNF as key upstream regulators of the effects of PRDX2KD (data not shown).

In subsequent experiments the effects of PRDX2KD on responses to electrical stimulation of contractions and exposure to H_2_O_2_ were examined in comparison with 2 control groups: unstimulated control (scramble shRNA -treated) myotubes and unstimulated PRDX2KD myotubes. Data obtained were comparable and hence comparison with unstimulated PRDX2KD myotubes are presented here for clarity.

### Effect of electrical stimulation of contractions on DEG in Scram treated myotubes

A total of 314 DEGs were identified in control Scram-treated myotubes following contractile activity compared with non-contracted Scram controls (Figure 3iA). The volcano plot in Figure 3iA demonstrates differential expression patterns, with upregulated genes highlighted in red and downregulated genes highlighted in blue. The bar graphs demonstrate the effect of contractions on relative changes in expressions of 17 selected key genes (Figure 3iB). Upregulated genes are shown in red with downregulated in blue. These show substantial relative increases in expression of genes involved in cellular energetics including *ATP6* and *COX1*, 2 and 3 sub-units of cytochrome oxidase compared with Scram control myotubes at rest. Down regulation of genes involved in extracellular matrix remodelling (including *COL3A1* and *COL5A2*) was also seen in the contracted myotubes compared with non-contracted myotubes. Using both KEGG (yellow) and Hallmark (orange) pathway analysis (Figure 3iC) there was a significant changes of a number of biologically relevant pathways associated with skeletal muscle responses to exercise, such as oxidative phosphorylation (adjusted p-value = 0.000615, odds ratio = 7.173), angiogenesis (adjusted p-value = 0.0154, odds ratio = 12.107) and mTOR signalling. The IPA analysis (Figure 3iD) indicated that the top canonical pathways affected by contractile activity were ribosomal RNA processing, oxidative phosphorylation and mitochondrial ATP production together with various aspects of the interferon-focussed inflammatory response.

### Effect of electrical stimulation of contractile activity on DEGs in PRDX2KD myotubes

Electrical stimulation of contracted PRDX2KD myotubes showed altered DEGs compared with non-contracted scramble-treated control myotubes (totalling 296 DEG genes). The volcano plot (Figure 3iiA) shows a different pattern of changes in DEG following contractions to those seen in the Scram control myotubes with a direct comparison of the 17 chosen genes highlighted in Figure 3iB. In contrast to the responses of Scram control myotubes, it is apparent that no significant changes in the expression of *COX2*, *COX3*, *TNNT2*, *COL3A1*, *COL5A2* and *LAMB1* were seen in the PRDX2KD myotubes following contractile activity. Thus, knock down of PRDX2 appears to have prevented the key changes in gene expression induced by electrically stimulated contractions that were seen in electrically stimulated Scram control myotubes (Figure 3i). The KEGG (yellow) and Hallmark (orange) pathway analysis (Figure 3iiC) also showed a substantially different set of key pathways activated by contractile activity in the PRDX2KD myotubes compared with non-contracted PRDXKD myotubes which illustrates the effect of PRDX2KD on myotube responses to contractile activity. IPA analysis (Figure 3iiD) also failed to identify any effect of the electrical stimulation on rRNA processing or oxidative phosphorylation. The DEG changes in several other pathways induced by contractile activity in the Scram-treated control myotubes were retained in the PRX2KD myotubes including modification of interferon signalling, stimulation of angiogenesis and changes in extracellular matrix organisation (Figure 3iiC & D).

**Figure 3(i).**
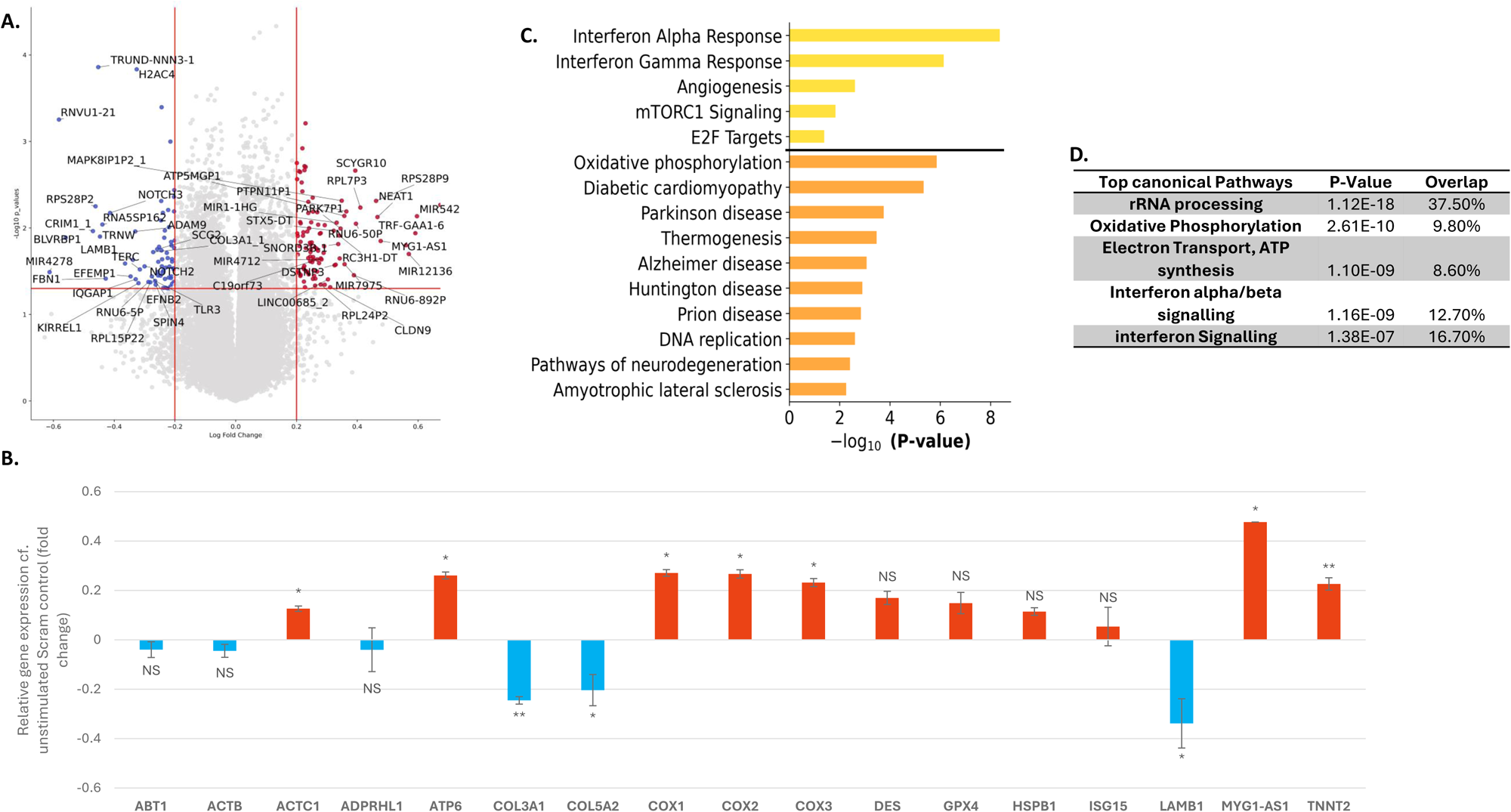
Effect of electrical stimulation of contractions on DEGs in control myotubes. (A) Volcano plot showing the DEGs between Scram control myotubes at rest and Scram myotubes that have undergone a 15 min period of electrically stimulated contractile activity. The x-axis indicates the log2 fold change, while the y-axis represents the -log10 (p-value). Genes with significant differential expression are highlighted in blue (downregulated) and red (upregulated). (B) Bar graphs showing the relative changes in expression of key genes discussed in the text. Upregulated genes shown in red with downregulated shown in blue. (C) KEGG (orange) and Hallmark (yellow) pathway analyses highlighting the pathways linked to the gene expression changes. (D) IPA analysis of the top five canonical pathways. Gene Key: ABT1 - Activator of Basal Transcription 1; ACTB - Actin Beta, ACTC1 - Actin Alpha Cardiac Muscle 1; ADPRHL1 - ADP-Ribosylhydrolase Like 1; ATP6 - ATP Synthase F0 Subunit 6; COL3A1 Collagen Type III Alpha 1 Chain; COl5A2 - Collagen Type V Alpha 2 Chain; COX1 - Cytochrome C Oxidase Subunit 1; COX2 - Cytochrome C Oxidase Subunit 2; COX3 - Cytochrome C Oxidase Subunit 3; DES – Desmin; GPX4 - Glutathione Peroxidase 4; HSPB1 - Heat Shock Protein Family B (Small) Member 1; ISG15 - Interferon-Stimulated Gene 15. LAMB1 - Laminin Subunit Beta 1; MYG1-AS1 - Myoglobin 1 Antisense RNA 1; TNNT2 - Troponin T Type 2 (Cardiac).

**Figure 3(ii).**
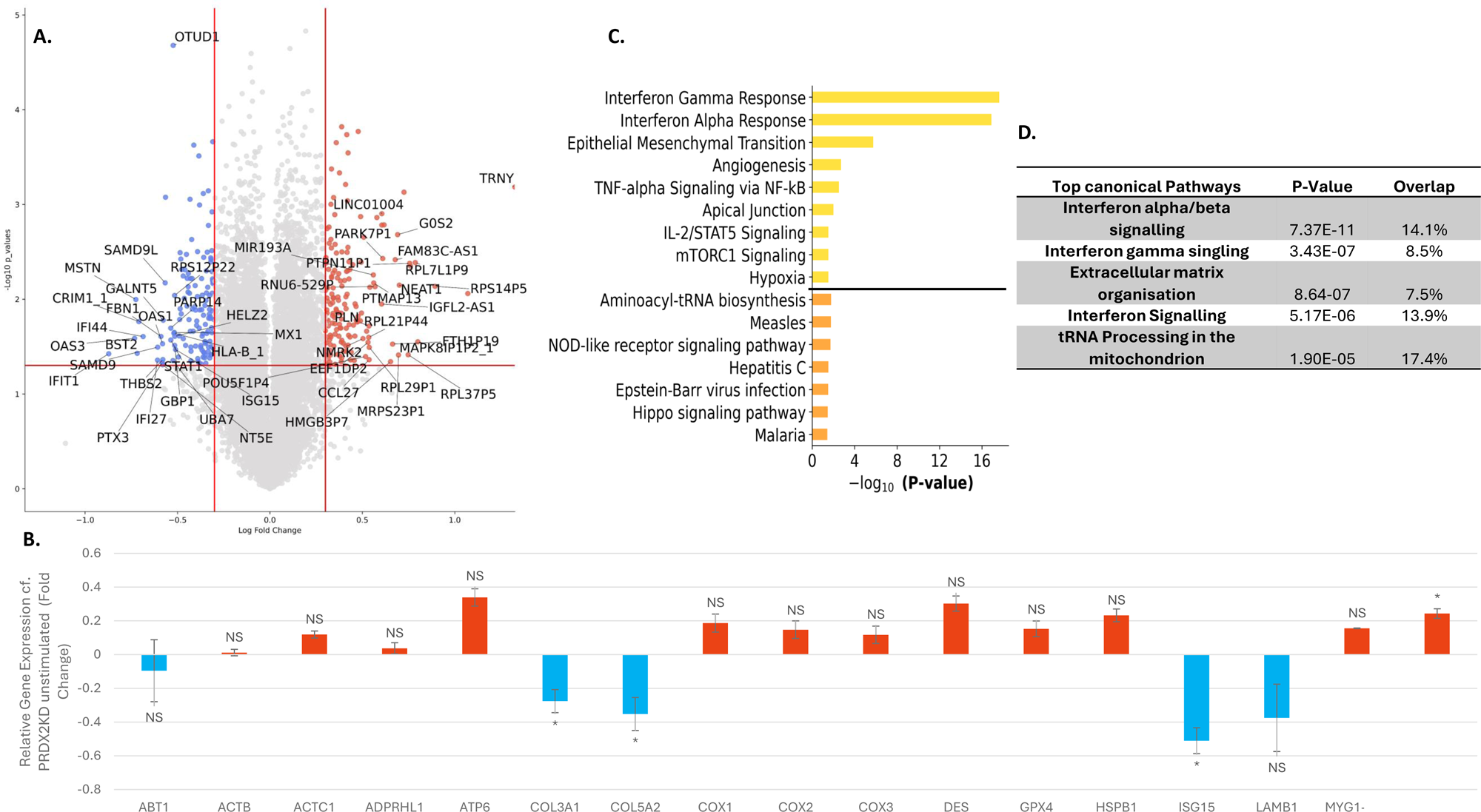
Effect of electrical stimulation of contractile activity on DEGs in PRDX2 KD myotubes. A) Volcano plot of the DEGs between PRDX2KD myotubes at rest and PRDX2 KD myotubes following 15 min electrical stimulation of contractions. The x-axis indicates the log2 fold change, while the y-axis represents the -log10 (p-value). Genes with significant differential expression are highlighted in blue (downregulated) and red (upregulated). (B) Bar graphs showing the relative changes in expression of key genes discussed in the text. Upregulated genes shown in red with downregulated shown in blue. (C) KEGG (orange) and Hallmark (yellow) pathway analyses highlighting the relationships between key genes linked to various pathways. (D) IPA analysis of the top five canonical pathways. Gene Key: as above.

### Effect of treatment with H_2_O_2_ on DEGs in Scram control myotubes

Analysis of the Scram control muscle myotubes following treatment with 5µM H_2_O_2_ showed 1180 DEGs (Figure 4iA) with many in common with the effects of electrical stimulation of contractions (shown in Figure 3iA). The same 17 genes (Figure 3) are again highlighted and are shown in Figure 4iB. These illustrate comparable upregulation in DEGs associated with oxidative phosphorylation and mitochondrial function (*ATP6*, *COX1*, *COX2*, *COX3*) also an upregulation in genes for redox sensitive proteins including HSPB1 (Figure 4iB). KEGG (yellow) and Hallmark (orange) pathway analysis outlined enrichment of a number of pathways indicating the extensive response of Scram control myotubes to 5µM H_2_O_2_ treatment (Figure 4iC). This included enrichment of Notch signalling and p53 and oxidative phosphorylation pathways. In parallel with the pattern seen following electrical stimulation of contractions, IPA analysis showed rRNA processing and electron transport/ATP synthesis in the top canonical pathways induced by 5µM H_2_O_2_ treatment.

### Effect of treatment with H_2_O_2_ on DEGs in PRDX2KD myotubes

Analysis of the PRDX2KD myotubes following 5µM H_2_O_2_ treatment revealed a blunted response compared with that seen in 5µM H_2_O_2_-treated Scram control myotubes (Figure 4iiA). Many of the individual genes differentially regulated by exposure to 5µM H_2_O_2_ in Scram control myotubes (Figure 4iB) showed no significant change, or in some instances showed a significant downregulation in the PRDX2KD myotubes following treatment with H_2_O_2_ including *COX1*, 2 and 3 and *ATP6* indicating a diminished mitochondrial response to H_2_O_2_ in PRDX2KO myotubes (Figure 4iiB). KEGG (yellow) and Hallmark (orange) pathway analysis identified several significant pathway changes induced by 5µM H_2_O_2_ treatment in the PRDX2KD myotubes. For example, TNF-alpha signalling which was identified in both KEGG and Hallmark analysis (Figure 4ii C). IPA analysis showed loss of the modification of rRNA processing and mitochondrial pathways previously seen in the H_2_O_2_ treated Scram control myotubes (Figure 4iD) with tRNA processing in mitochondria being retained as a key response in both control and KD myotubes (Figure 4iiD).

In a separate experiment Scram control myotubes and PRDX2KD myotubes were treated with a lower concentration of H_2_O_2_ (2.5µM). These data showed many of the same changes in DEGs as were seen with 5µM H_2_O_2_ including upregulation of *COX* genes in the Scram control myotubes that were not seen in the PRDX2KD myotubes, but the pathway analyses did not identify these as major pathways. These data are presented as supplementary data for completeness (Supplementary Figures S1 and S2).

### Comparison of DEG induced by electrical stimulation of contraction or exposure to 5µM H_2_O_2_ in Scram control or PRDX2KD myotubes

Figure 5A shows a Venn diagram of the number of DEGs that are common or specific to the electrically stimulated or H_2_O_2_-treated groups of Scram control myotubes. The data indicate 34 common DEGs between the 5µM H_2_O_2_ treated and the contracted groups of Scram control myotubes. Figure 5B shows the IPA analysis of these common genes and indicates that they focus on rRNA processing, oxidative phosphorylation and sirtuin signalling as anticipated from the data shown above. A list of the common DEGs is presented in Table 1 showing that 33 of 34 the DEGs moved in the same direction in the 2 different treatment groups with 29 upregulated and 4 downregulated. The remaining DEG (*RNU6-5P*) was upregulated following contractions and downregulated in myotubes treated with H_2_O_2_. Table 1 also shows which DEGs contribute to the pathway identification in the IPA analysis. A comparison Venn diagram showing the common DEGs between PRDX2KD myotubes following either electrical stimulation or treatment with 5µM H_2_O_2_ is shown in Figure 5C with the IPA analysis of the 15 shared genes shown in Figure 5D. A list of common DEGs between these conditions in the PRDXKD myotubes is presented in Table 2. As anticipated, none of the DEGs overlap with the common responses to contractions and 5µM H_2_O_2_ in Scram control myotubes. Of the 15 common DEGs in the PRDX2KD myotubes, 14 were up regulated. IPA analysis of the common DEGs in the PRDX2KD myotubes identified modification of tRNA processing in the mitochondria as a key common pathway modified by electrical stimulation or peroxide treatment.

**Table 1.**
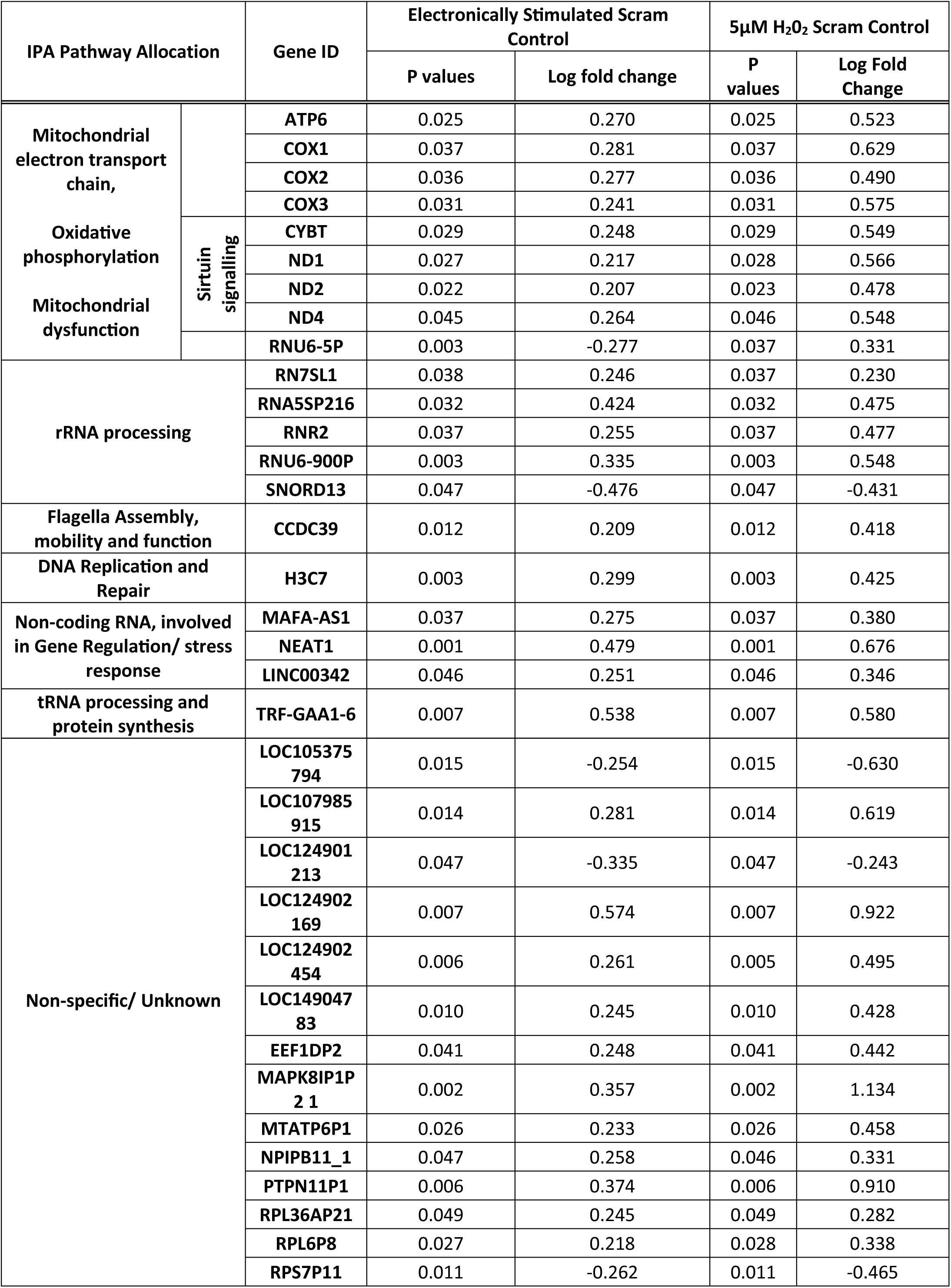
Common DEGs between Scram control myotubes following 15 minutes of contractile activity or exposure to 5µM H_2_O_2_ showing log fold change and P value for significance of the change. Those genes contributing to the pathways identified by IPA analysis are also indicated.

**Table 2.**
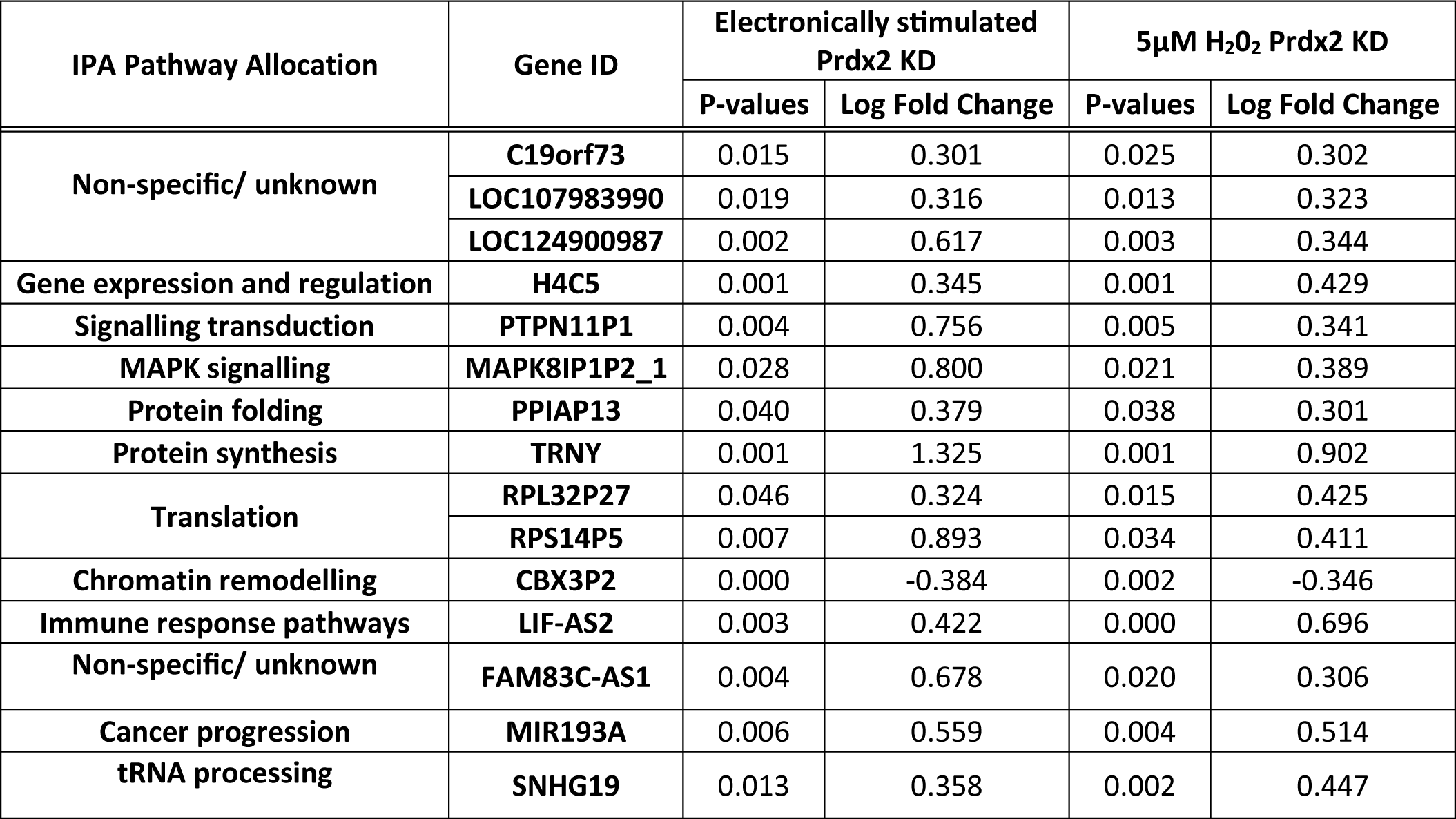
Common DEGs between PRDX2KD myotubes following 15 minutes of contractile activity or exposure to 5µM H_2_O_2_ showing log fold change and P value for significance of the change. Those genes contributing to the pathways identified by IPA analysis are also indicated.

| IPA Pathway Allocation | Gene ID | Electronically stimulated Prdx2 KD |  | 5µM H<sub>2</sub>O<sub>2</sub> Prdx2 KD |  |
| --- | --- | --- | --- | --- | --- |
|  |  | P-values | Log Fold Change | P-values | Log Fold Change |
| Non-specific/ unknown | **C19orf73** | 0.015 | 0.301 | 0.025 | 0.302 |
|  | **LOC107983990** | 0.019 | 0.316 | 0.013 | 0.323 |
|  | **LOC124900987** | 0.002 | 0.617 | 0.003 | 0.344 |
| Gene expression and regulation | **H4C5** | 0.001 | 0.345 | 0.001 | 0.429 |
| Signalling transduction | **PTPN11P1** | 0.004 | 0.756 | 0.005 | 0.341 |
| MAPK signalling | **MAPK8IP1P2_1** | 0.028 | 0.800 | 0.021 | 0.389 |
| Protein folding | **PPIAP13** | 0.040 | 0.379 | 0.038 | 0.301 |
| Protein synthesis | **TRNY** | 0.001 | 1.325 | 0.001 | 0.902 |
| Translation | **RPL32P27** | 0.046 | 0.324 | 0.015 | 0.425 |
|  | **RPS14P5** | 0.007 | 0.893 | 0.034 | 0.411 |
| Chromatin remodelling | **CBX3P2** | 0.000 | -0.384 | 0.002 | -0.346 |
| Immune response pathways | **LIF-AS2** | 0.003 | 0.422 | 0.000 | 0.696 |
| Non-specific/ unknown | **FAM83C-AS1** | 0.004 | 0.678 | 0.020 | 0.306 |
| Cancer progression | **MIR193A** | 0.006 | 0.559 | 0.004 | 0.514 |
| tRNA processing | **SNHG19** | 0.013 | 0.358 | 0.002 | 0.447 |

## Discussion

This work examined the role of PRDX2 in responses of human skeletal muscle to contractile activity or exposure to low physiological concentrations of H_2_O_2_. The human skeletal muscle cell line readily differentiates into multinucleated myotubes that are contractile when activated by field electrical stimulation. These myotubes showed a robust response to electrically stimulated contractions with a spectrum of transcriptional alterations that are also seen in the responses of skeletal muscle to aerobic contractile activity in vivo [40] (Figure 3), these included upregulation of angiogenesis pathways, oxidative phosphorylation, rRNA processing and inflammatory responses. These responses were compared with the DEGs induced by exposure of the myotubes to low concentrations of H_2_O_2_ which our previous studies have shown induce oxidation and dimerization of PRDX2 in skeletal muscle fibres without significant formation of hyper-oxidised PRDX [26]. We reasoned that comparison of the effects of exposure to contractile activity and “physiological” levels of H_2_O_2_ in control myotubes and myotubes lacking PRDX2 would allow identification of DEGs and enriched pathways that are stimulated by contractions, redox-regulated and mediated by oxidation of PRDX2.

### Knock down of PRDX2 in human skeletal muscle myotubes

The shRNA strategy to knock down PRDX2 in myoblasts which were then fused into multi nuclear myotubes was successful with good uptake of the AAV vector as illustrated by the red MCherry tag integrated into the plasmid (Figure 1). This led to an efficient knockdown assessed using rt-qPCR with a reduction of 98% in PRDX2 RNA levels at ten days after of transduction (Figure 1). There was no difference in the PRDX2 mRNA content of the myotubes between the five and ten-day time points post-transduction suggesting that maximal RNA knockdown was achieved within this timeframe. Western blot analysis showed a significant decrease in PRDX2 content after five days of transduction an average decrease of 44.8% in PRDX2 content after 10 days of transduction (Figure 1). These data are in accordance with a relatively slow turnover of the PRDX2 protein [42].

### Effect of knock down of PRDX2 on differentially expressed genes (DEGs) in human skeletal muscle myotubes at rest

RNA sequencing of the myotubes at ten days post-KD of PRDX2 compared with control myotubes treated with scrambled shRNA showed many DEGs (Figure 2) and illustrates the extensive role played by PRDX2 in homeostatic pathways in skeletal muscle. Markedly influenced by Prdxd2KD were DEGs for proteins involved in energy metabolism and mitochondrial function. Examples included *COX5A, COX5B, COX6B1, COX7A1, COX7B, COX7C, COX8A* (Log fold change = -0.4257, P value = 0.0027, Log fold change = -0.4344, P value = 0.0027, Log fold change = -0.3090, P value = 0.0044, Log fold change = -0.4764, P value = 0.0038, Log fold change = -0.3855, P value = 0.0085, Log fold change = -0.3233, P value = 0.0094, Log fold change = -0.3355, P value = 0.0043, Figure 2) which were all downregulated indicating that PRDX2 may play a significant role in regulating mitochondrial function and energy metabolism in muscle myotubes at rest. The downregulation of genes encoding subunits of *COX* implies potential impairment in mitochondrial respiration and energy production. *COX1* was upregulated (Log fold change = 0.4748 P value= 0.00083) potentially suggesting a compensatory response to the dysregulation of other *COX* subunits. Pathway analysis of the DEG data by the KEGG and Hallmark programmes highlighted changes in oxidative phosphorylation as the major affected pathway which was confirmed by IPA analysis which also identified Peroxisome proliferator-activated receptor gamma coactivator 1-alpha (PPARCIA) as a major regulator affected by the KD potentially reflecting an inhibitory effect on mitochondrial biogenesis in the PRDX2KD myotubes.

In addition to the metabolic alterations, PRDX2KD induced a change in a number of DEGs that have been associated with the regulation of cellular structure and the cytoskeleton including a down regulation of *ACTA1* (Log fold change = -0.4621 p value = 0.01709). *ACTA1* is a key protein involved in the structure and function of skeletal muscle fibres, specifically in the regulation of muscle contraction. Its downregulation following PRDX2 knockdown suggests potential disruptions in the organization and function of the cytoskeleton in muscle myotubes. The cytoskeleton plays crucial roles in maintaining cell shape, providing structural support, and facilitating intracellular transport and signalling.

These data also showed an overall downregulation of genes associated with the inflammatory response following PRDX2KD. Decreases in genes such as *CCL20, CCL27, CXCL2*, and *EGLN1*, suggest that there may be a potential dampening effect of chemokines produced by muscle. IPA analysis also identified a major effect on *TGFβ1* and *TNF* regulation (data not shown). Additionally, PRDX2 knockdown positively influenced the expression of genes involved in apoptosis regulation (*BBC3, GADD45A*) and cytokine signalling (*IL11, INHBA, LIF*).

### Effect of electrical stimulation of contractions on DEG in Scram control myotubes

Comparison between resting and electrically stimulated skeletal muscle myotubes showed a spectrum of transcriptional alterations which indicated that this human myotube model shows multiple aspects of the responses to aerobic contractile activity that are also seen in mature skeletal muscle in vivo [46–48]. Pathway analysis (Figure 3) indicated that the contractile activity induced changes in angiogenesis and oxidative phosphorylation/electron transport with increases in ribosomal RNA processing and interferon signalling, all of which are acknowledged responses to contractile activity in muscle in vivo [40].

Prominent among the DEGs observed in the contracted myotubes were multiple genes involved in mitochondrial function. These included *ATP6* (log fold change = 0.267, p-value = 0.017) which is a pivotal player in cellular energetics and encodes a subunit of ATP synthase (Figure 3i). This was associated with upregulation of *COX1* (log fold change = 0.278, p-value = 0.018), *COX2* (log fold change = 0.274, p-value = 0.018), and *COX3* (log fold change = 0.238, p-value = 0.032) which encode subunits of cytochrome c oxidase, a key enzyme complex in the mitochondrial electron transport chain. Data obtained here are limited to changes in mRNA expression since information on effects on relevant protein contents and mitochondrial function were not obtained, but we infer that the upregulation of these genes indicates a stimulus to enhance mitochondrial respiration and oxidative phosphorylation in response to contractions, facilitating ATP production.

Other changes induced by the electrically stimulated contractions include upregulation of *MYG1*-*AS1* (log fold change = 0.485, p-value = 0.014) a long non-coding RNA (lncRNA), implicated in regulating gene expression patterns underlying muscle development and function. A significant upregulation of the DEGs for *TNNT2* (log fold change = 0.231, p-value = 0.002) was also observed. This gene encodes troponin T, a crucial component of the troponin complex governing muscle contraction regulation.

In contrast there were several downregulated DEGs associated with extracellular matrix (*ECM*) remodelling and structural including *COL3A1* (log fold change = -0.250, p-value = 0.004) and COL5A2 (log fold change = -0.207, p-value = 0.042), encoding collagen proteins crucial for ECM organization. This downregulation may signify a remodelling of the ECM to accommodate the dynamic mechanical stresses imposed by contractile activity.

### Effect of electrical stimulation of contractile activity on DEGs in PRDX2KD myotubes

Electrical stimulation of contractile activity in PRDX2KD myotubes led to a substantially different set of DEGs compared with contracted Scram control myotubes, most notably there was no significant up-regulation of *COX1, COX2,* and *COX3* gene expression and pathway analysis failed to identify upregulation of ribosomal RNA processing, oxidative phosphorylation or electron transport in these myotubes (Figure 3ii). As anticipated, a number of pathways modified by contractile activity in the control myotubes were also modified in the PRDX2KD myotubes including interferon signalling, angiogenesis and extracellular matrix remodelling. The abolished of these responses to contractile activity in the PRDX2KD myotubes illustrates the specificity of the potential roles played by PRDX2 in muscle responses during exercise. These include important responses such as interferon signalling which is important for immune modulation, muscle inflammation, atrophy, and impaired regeneration [49].

Changes in extracellular matrix organisation occurred following contractile activity in both Scram control and PRDX2KD myotubes following stimulation to contract. This included a significant downregulation of key genes involved in extracellular matrix (ECM) organization, The downregulation of ECM genes suggests alterations in the genes responsible for control of muscle architecture and mechanical properties, which are indispensable for efficient muscle function during exercise are not regulated by PRDX2. tRNA processing in mitochondria was also indicated by IPA analysis as a key response to contractions that was unaffected by PRDX2KD.

### Effect of treatment with H_2_O_2_ on DEGs in Scram control myotubes

Two concentrations of H_2_O_2_ were examined, 2.5 and 5µM H_2_O_2_. These concentrations were chosen because of a lack of specific knowledge on the concentration of extracellular H_2_O_2_ that would be required to lead to a relevant “physiological” increase in cytosolic H_2_O_2_. While it is acknowledged that cytosolic H_2_O_2_ concentrations approximate to 10nM at rest [50], there is considerable debate about how much this will increase following addition of micromolar concentrations of extracellular H_2_O_2_, due to differing views on the concentration drop that occur across the plasma membrane [16, 51]. In previous studies 2.5 - 5µM H_2_O_2_ was found to induce oxidation of PRDX2 in isolated muscle fibres without significant hyper-oxidation of PRDXs [26] and hence these concentrations were chosen for use here. Treatment with 5µM H_2_O_2_ induced clear changes in DEGs from Scram control myotubes that in multiple ways paralleled the changes seen with electrical stimulation of contractions. Exposure of control myotubes to 2.5µM H_2_O_2_ generally showed similar patters to 5uM H_2_O_2_ with, for instance, upregulation of genes associated with oxidative phosphorylation, but the pathway analyses were less definitive, and these data are provided in Supplementary data (Figure S1) for completeness.

Multiple DEGs were identified in response to H_2_O_2_ treatment of Scram control myotubes (Figure 4) and these included substantial overlap with the DEGs induced by contractile activity (Figure 3). As with data for contracted Scram control myotubes the top canonical pathways identified by IPA analysis included ribosomal RNA processing and oxidative phosphorylation/electron transport/ATP synthesis with oxidative phosphorylation also identified by the KEG and hallmark programmes. Specifically ATP-related genes, including *ATP6* (log fold change = 0.517, p-value = 0.025) and *ATP8* (log fold change = 0.506, p-value = 0.029) and *COX1*, 2 and 3 (log fold change = 0.621, p-value = 0.037, log fold change = 0.484, p-value = 0.035, log fold change = 0.56, p-value = 0.031) were all significantly up regulated following treatment with 5µM H_2_O_2_ (Figure 4iA & B). Numerous genes associated with mitochondrial complex I, NADH dehydrogenase where also upregulated including ND1 (log fold change = 0.559, p value = 0.027), *ND2* (log fold change = 0.472, P value = 0.022), *ND4* (log fold change = 0.541, P value = 0.045), and *ND6* (log fold change = 0.426, P value = 2.3E-05, Figure 5i). These multiple gene changes all support a coordinated response aimed at augmenting electron transport and ATP synthesis to meet increased energy demands.

**Figure 4(i).**
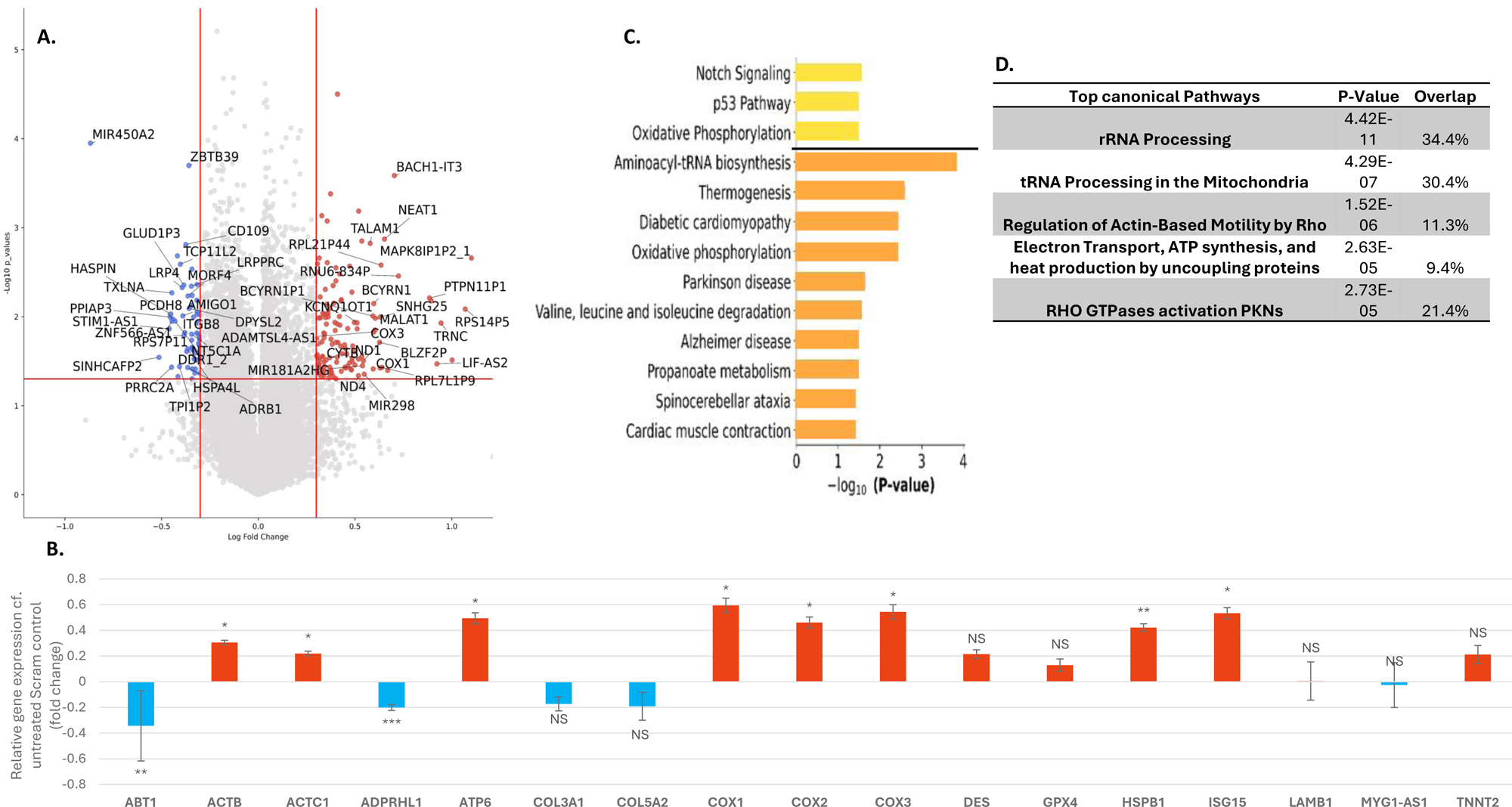
Effect of treatment with 5µM H_2_O_2_ on DEGs in scram control myotubes. (A) Volcano plot showing the distribution of the DEGs between Scram control untreated and 5µM H_2_O_2_-treated Scram myotubes. The x-axis indicates the relative log2 fold change, while the y-axis represents the -log10 (p- value). Genes with significant differential expression are highlighted in blue (downregulated) and red (upregulated). (B) Bar graphs showing the relative changes in expression of key genes discussed in the text. Upregulated genes shown in red with downregulated shown in red. (C) KEGG (orange) and Hallmark (yellow) pathway analyses highlighting the pathways linked to the gene expression changes (D) IPA analysis of the top five canonical pathways. Gene Key: as in Figure 3.

**Figure 4(ii).**
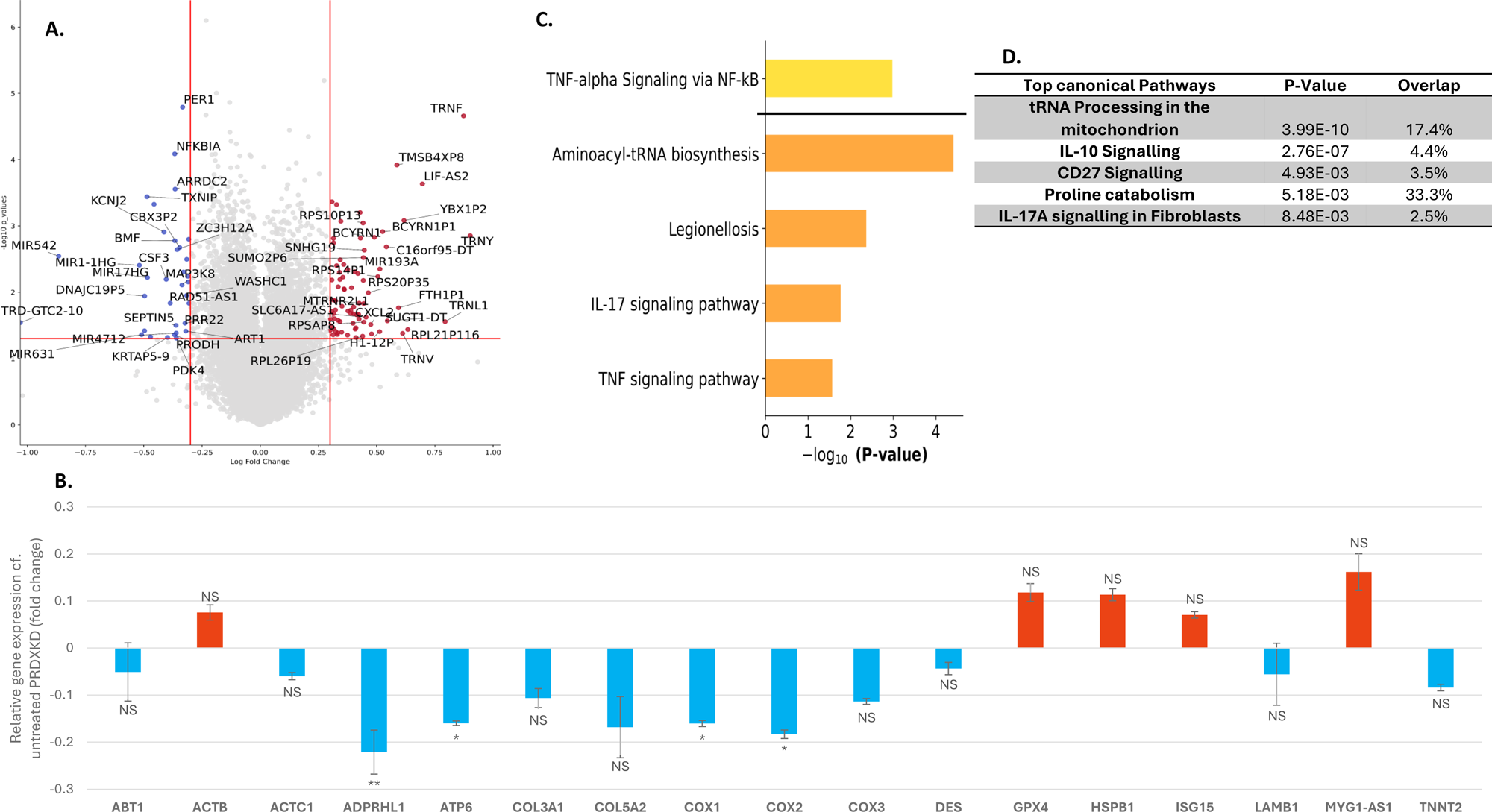
Effect of treatment with 5µM H_2_O_2_ on DEGs in PRDX2KD myotubes. (A) Volcano plot showing the DEGs between untreated PRDX2KD myotubes and PRDX2KD myotubes treated with 5µM H_2_O_2_. The x-axis indicates the relative log2 fold change, while the y-axis represents the -log10 (p-value). Genes with significant differential expression are highlighted in blue (downregulated) and red (upregulated). (B) Bar graphs showing the relative changes in expression of key genes discussed in the text. Upregulated genes shown in red with downregulated shown in blue. (C) KEGG (orange) and Hallmark (yellow) pathway analyses highlighting the pathways linked to the gene expression changes. (D) IPA analysis of the top five canonical pathways. Gene Key: as above

A number of genes associated with muscle structure and function were also upregulated by the H_2_O_2_ exposure, including *ACTB* (log fold change = 0.319, p-value = 0.022), *ACTC1* (log fold change = 0.234, p-value = 0.031), and *ACTG1P15* (log fold change = 0.230, p-value = 0.044) upregulated. These genes encode actin isoforms, essential components of the cytoskeleton involved in muscle contraction (Figure 4i). In contrast several genes implicated in negative regulation of muscle function showed downregulation following H_2_O_2_ treatment. For instance, *ABT1* (log fold change = -0.354, p-value = 0.007) and *ADPRHL1* (log fold change = -0.207, p-value = 0.001) are downregulated. These genes are involved in processes such as protein synthesis and DNA repair. The peroxide exposure also modified the expression of redox-sensitive genes involvedin cell stress responses and antioxidant defence mechanisms, such as *HSPB1* (log fold change = 0.439, p-value = 0.003) and *ISG15* (log fold change = 0.552, p-value = 0.048), is significantly altered following H_2_O_2_ treatment.

### Effect of treatment with H_2_O_2_ on DEGs in PRDX2KD myotubes

PRDX2KD had a substantial effect on responses to treatment with 5µM H_2_O_2_ and in a similar manner to those post-contractions, multiple genes associated with mitochondrial function and oxidative phosphorylation which were upregulated post-contractions in the Scram control myotubes (including *COX1*, *COX2*, and *COX3*) were either unchanged or down regulated following peroxide treatment in the PRDX2KD myotubes. This pattern was also found for multiple mitochondrial complex I genes (*ND1*, *ND2*, *ND4*, *ND6*, *ATP 6* and *ATP 8*), none of which were upregulated in the PRDX2 myotubes (data not shown in detail in Figure 4). These findings reiterate the finding from other studies implicating PRDX2 as an important regulatory molecule in expression of some of the key mitochondrial genes involved in oxidative phosphorylation and ATP synthesis [27, 28].

In common with the effects of contractile activity, PRDX2KD myotubes exposed to 5µM H_2_O_2_ retained some of the pathway activation seen in Scram control myotubes in response to H_2_O_2_ with the IPA Pathway analysis identifying changes in tRNA processing in mitochondria and activation of inflammatory pathways as key retained pathways stimulated by H_2_O_2._

Treatment of myotubes with 2.5µM H_2_O_2_ (Supplementary data, Figure S1) also led to an increase in DEGs of multiple genes included in energy metabolism in Scram control myotubes in common with the findings from electrically stimulated contraction data. Expression levels of the genes for *ATP6* (log fold change = 0.445, p-value = 0.041), *COX2* (log fold change = 0.48, p-value = 0.039), and *COX3* (log fold change = 0.489, p-value = 0.028) were elevated in Scram control myotubes following treatment with H_2_O_2_ and additionally there was an increase in the expression of *ND3* and *ND6* genes (Log fold change = 0.3637, P value = 0.04298; Log fold change = 0.348129272, P value = 0.01680). *ND3* and *ND6* genes encode subunits essential for the functionality of the mitochondrial electron transport chain complex NADH dehydrogenase. Despite these changes IPA pathway analysis did not robustly identify oxidative phosphorylation or mitochondrial changes as key pathways affected by this level of peroxide, although oxidative phosphorylation was highlighted in the KEG analysis (Supplementary data Figure S1). In a similar pattern to that seen with the 5µM H_2_O_2_ treatment, the changes in mitochondrial genes seen in Scram control myotubes following treatment with 2.5µM H_2_O_2_ were abolished in the PRDX2KD myotubes (Supplementary data, Figure S2), although again some pathways (e.g Hedgehog signalling) were more robustly modified by 2.5µM H_2_O_2_ treatment in both Scam control and PRDX2KD myotubes.

Thus, the data from these experiments show that electrical stimulation of contractions induces multiple changes in the DEGs in myotubes and that key pathways are activated to increase oxidative phosphorylation and ribosomal RNA processing. These changes were mimicked by exposure of myotubes to 5µM H_2_O_2_ and these effects of both stresses were effectively abolished by knockdown of PRDX2. We therefore conclude that specific gene responses in human muscle myotubes to increase oxidative phosphorylation and ribosomal RNA (rRNA) processing in response to contractile activity are likely to be redox-regulated and mediated by PRDX2.

### Comparison of DEGs induced by electrical stimulation of contractions and exposure to 5µM H_2_O_2_ in Scram control and PRDX2KD myotubes

Figure 5 shows a Venn diagram illustrating the overlap between DEG induced by electrical stimulation and treatment with 5µM H_2_O_2_ in Scram control (Figure 5A) and PRDX2KD (Figure 5C) myotubes. In Scram control myotubes 34 DEGs were found in common between the 2 different treatment groups (Contractions or H_2_O_2_) with 33 changing in the same direction (Table 1) with 29 upregulated and 4 downregulated. The remaining DEG (*RNU6-5P*) was downregulated following contractions and upregulated in myotubes treated with H_2_O_2_. IPA analysis of these genes identified ribosomal RNA processing, oxidative phosphorylation and sirtuin signalling as the key pathways involved (Figure 5B). None of these DEGs were overlapping with the common genes seen in PRDX2KD cells following electrical stimulation and treatment with 5µM H_2_O_2_. In PRDX2KD myotubes 15 DEG were shared between the electrically stimulated and peroxide treated cells with 14 being upregulated and 1 down regulated (Table 2). Pathway analysis of the common DEGs in the PRDX2KD myotubes yielded little information with no overlap with the pathways identified in the Scram control myotubes although modification of tRNA processing in mitochondria was a major response to peroxide treatment that was maintained in the absence of PRDX2.

**Figure 5.**
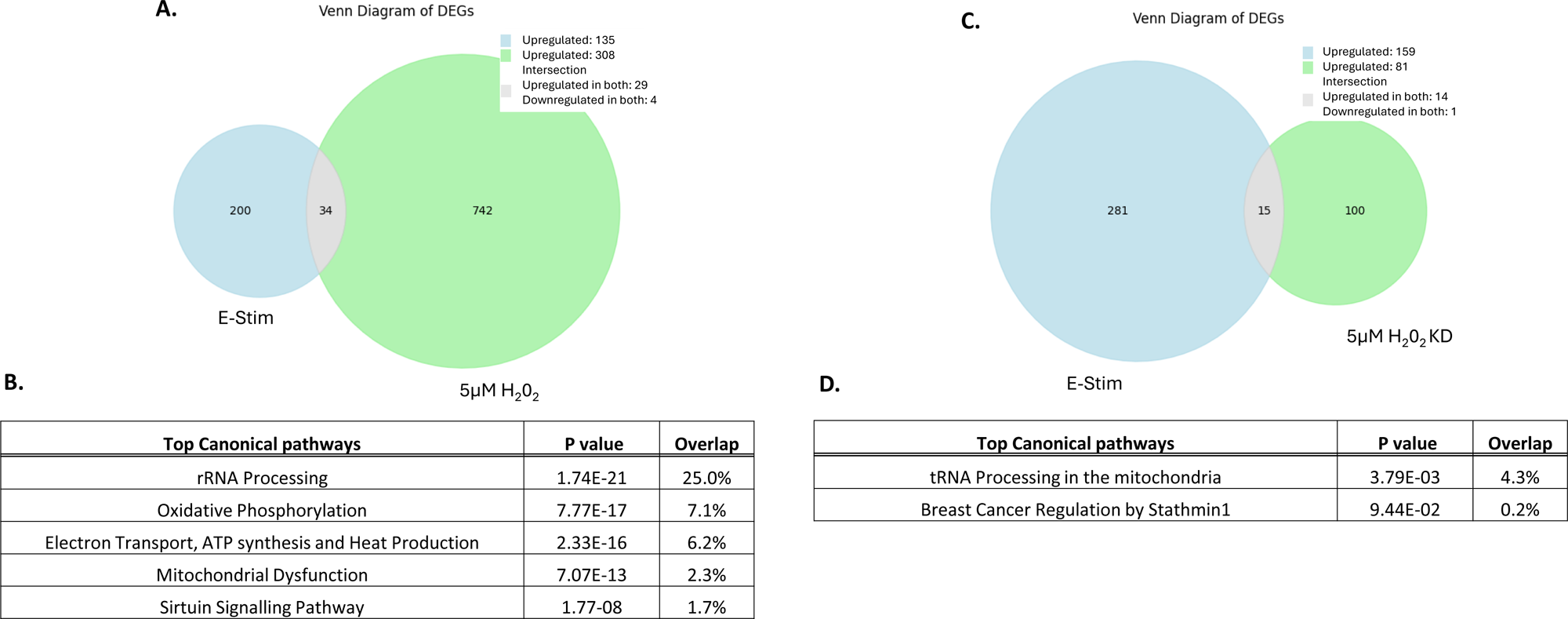
Comparison of DEG induced by electrical stimulation of contraction and exposure to 5µM H2O2 in PRDX2KD myotubes. (A) Venn diagram of DEGs following treatment with 5µM H2O2 or electrical stimulation of contractions in Scram control myotubes. 34 common genes were identified between the 2 conditions and IPA analysis of these genes indicated that they were involved in rRNA processing, mitochondrial function, oxidative phosphorylation and sirtuin signalling (Figure 5B). Figure 5C shows the Venn diagram of DEGs following treatment with 5µM H2O2, or electrical stimulation of contractions in PRDX2KD myotubes. 15 common genes were identified between the 2 conditions with IPA analysis identifying only 2 pathways due to the low number of common genes, with tRNA processing in the mitochondria and breast cancer regulation by stathmin1 being identified as the prominent common pathways.

It is relevant to speculate why these 15 genes are commonly expressed in contracted and peroxide-treated PRDX2KD myotubes, but not detected as a common response to contractions in Scram control myotubes. PRDX2 has a major role as the key regulator of cytosolic H_2_O_2_ in cells and acts as an antioxidant preventing cell damage [53]. In the absence of the enzyme it is likely that cytosolic H_2_O_2_ will increase to a greater extent following electrical stimulation or exposure to 5µM H_2_O_2_ than in control myotubes potentially leading to oxidative damage and activation of an alternative set of cellular stress responses [53]. Such genes may include tRNA processing in mitochondria as appears to have occurred here. Alternatively PRDX have roles as protein chaperones [54] and a lack of this activity might also lead to activation of alternative stress response pathways in response to contractile activity or peroxide treatment.

In conclusion, this study has revealed substantial commonality in the DEGs induced by exposure of human skeletal muscle myotubes to a 15 min period of electrically-stimulated isometric contractions compared with 5μM H_2_O_2_ exposure. Common DEGs in between exposure to H2O2 and electrical stimulation of contractions included upregulation of genes associated with increased mitochondrial oxidative phosphorylation, including *COX1, COX2, COX3* and *ATP6* and upregulation of genes associated with ribosomal RNA processing. Upregulation of mitochondrial capacity including oxidative phosphorylation is a key response of muscle to contractions and exercise leading to increased exercise capacity. Previous data for studies using high dose nutritional antioxidants has suggested that increased ROS in exercising muscle stimulate mitochondrial biogenesis [55] and Xia et al have recently shown that deletion of PRDX2 in *C.elegans* leads to a reduction in mitochondrial function and decreased adaptations following swimming exercise [27]. Here we have shown that KD of PRDX2 in human muscle myotubes prevented the upregulation of genes associated with increased oxidative phosphorylation and mitochondrial ATP production that occurs in response to contractile activity.

Data presented also indicate that contraction-induced changes in the processing of muscle ribosomal RNA may be redox-regulated and mediated by PRDX2. Changes in the processing or rRNA is a key component of the adaptation of cells to stress to maintain nucleolar integrity [56]. Skerlovsky and colleagues [57] have speculated that ROS act as signal transducers to promote endonucleocytic cleavage of 25S ribosomal RNA, a putative regulatory region on the surface of the 60S ribosomal subunit. They speculate that this enables cells to adapt to successfully counteract stress. The process does not appear to have been studied in muscle cells, but our data suggest that similar stress responses may occur in skeletal muscle in response to electrical stimulation or peroxide exposure and that these are mediated by PRDX2.

Although our data appear to support the concept that PRDX2 acts as a medicator of at least these 2 redox-regulated responses to contractile activity, these represent only a very small part of the potential responses to exercise that may be redox-regulated. In addition, PRDX1 and 3 are also rapidly oxidised in response to contractile activity in muscle [26] and might potentially play an analogous role in mediating other adaptations to contractile activity. The approach followed here may additionally provide a route to identify such effects.

## Acknowledgments

The authors would like to acknowledge the generous financial support from UKRI Medical Research Council (grant number MR/V03412X/1) and the University of Liverpool core Central Genomic Facility who undertook the RNA sequencing analysis.

## Contributions

RAH, STMB, CAS conducted the laboratory work, developed the RNAseq and statistical analysis and contributed to the writing of the paper, VM developed the immortalised human muscle cell line and SWJ established this model in the Liverpool laboratory, and both contributed to the paper. MJJ and AMcA devised the project, obtained the funding, supervised the project and contributed to data analysis, interpretation and writing of the manuscript.

## Supplementary data

**Figure S1.**
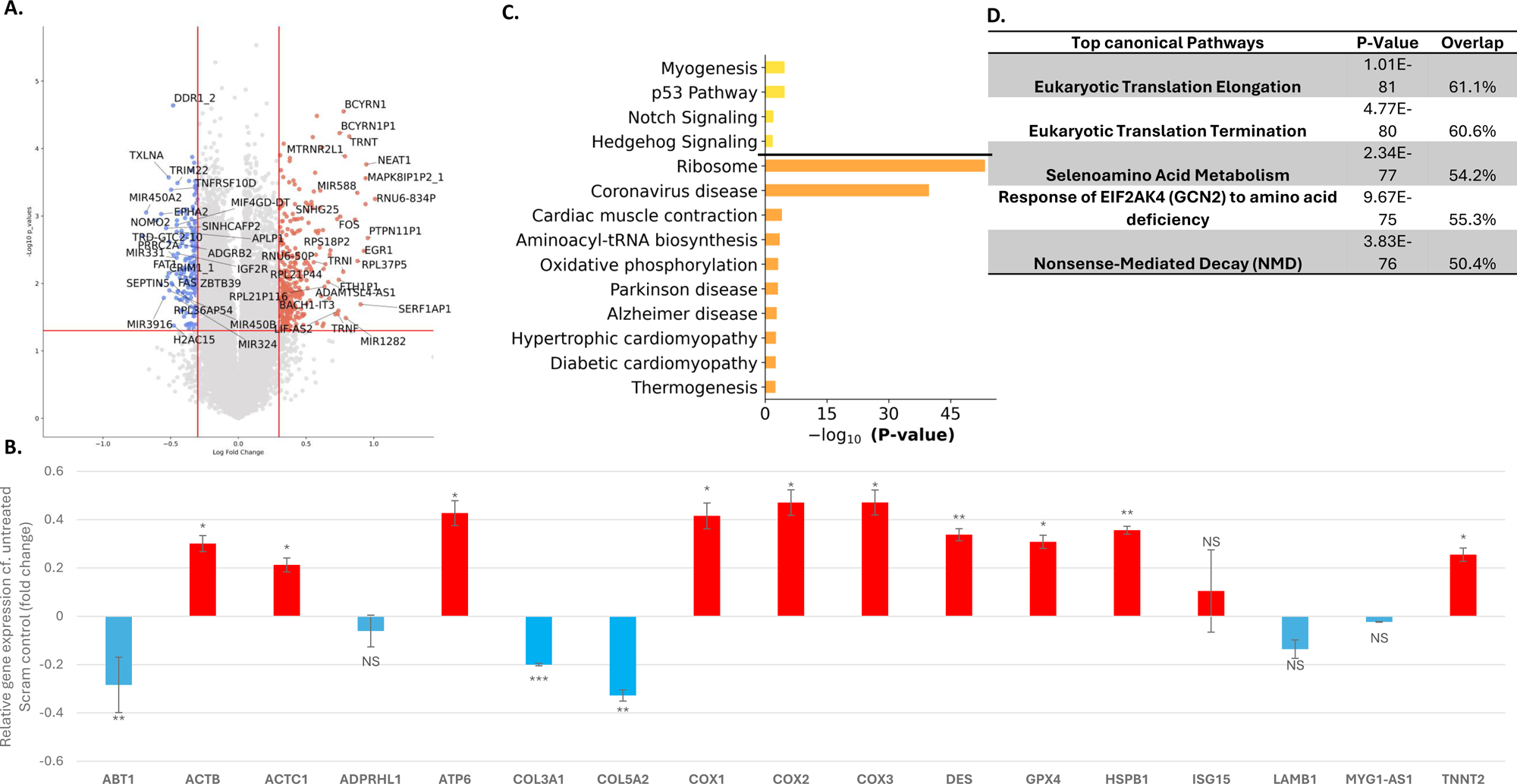
Effect of treatment with 2.5µM H_2_O_2_ on DEGs in Scram control myotubes. (A) Volcano plot highlighting differentially expressed genes, the x-axis indicates the relative log2 fold change, while the y-axis represents the -log10 (p-value). Genes with significant differential expression are highlighted in red (upregulated) and blue (downregulated). B) Bar graphs showing the relative changes in expression of key genes discussed in the text. Upregulated genes shown in red with downregulated shown in blue. (C) KEGG (orange) and Hallmark (yellow) pathway analyses highlighting the pathways linked to the gene expression changes. (D) The top 5 Canonical pathways identified from the IPA analysis.

**Figure S2.**
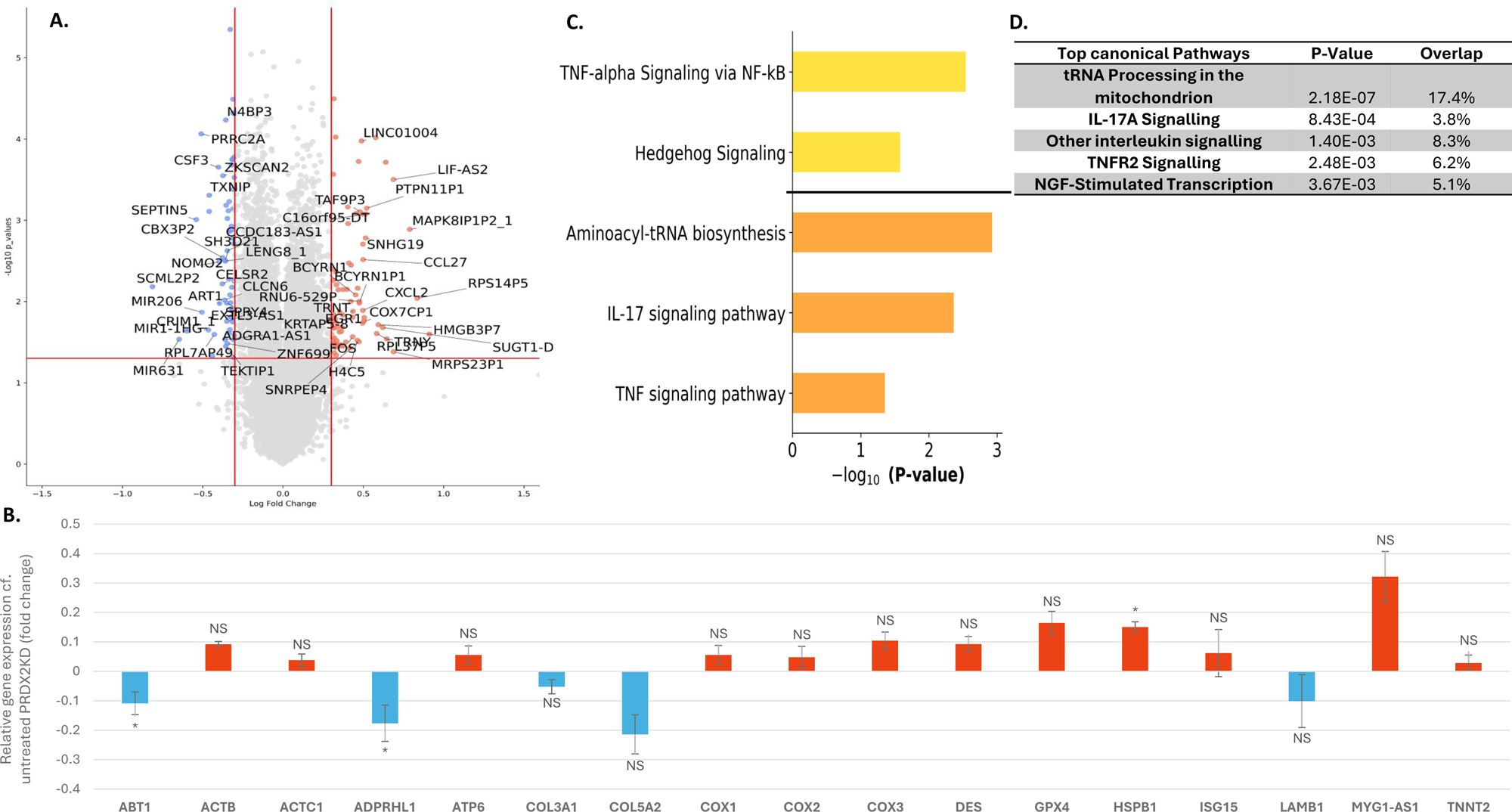
Effect of treatment with 2.5µM H_2_O_2_ on DEGs in PRDX2 KD myotubes. (A) Volcano plot showing the differentially expressed genes between untreated PRDX2KD myotubes and PRDX2 KD myotubes treated with 2.5µM H_2_O_2_. The x-axis indicates the relative log2 fold change, while the y-axis represents the -log10 (p-value). Genes with significant differential expression are highlighted in red (upregulated) and blue (downregulated). (B) Bar graphs showing the relative changes in expression of key genes discussed in the text. Upregulated genes shown in red with downregulated shown in blue. (C) KEGG (orange) and Hallmark (yellow) pathway analyses highlighting the pathways linked to the gene expression changes. (D) The top 5 Canonical pathways identified from the IPA analysis.

## References

[1] G.L. Close, A.C. Kayani, T. Ashton, A. McArdle, M.J. Jackson, Release of superoxide from skeletal muscle of adult and old mice: an experimental test of the reductive hotspot hypothesis, Aging Cell 6(2) (2007) 189–95.

[2] J. Palomero, D. Pye, T. Kabayo, D.G. Spiller, M.J. Jackson, In situ detection and measurement of intracellular reactive oxygen species in single isolated mature skeletal muscle fibers by real time fluorescence microscopy, Antioxid Redox Signal 10(8) (2008) 1463–74.

[3] S.K. Powers, M.J. Jackson, Exercise-induced oxidative stress: cellular mechanisms and impact on muscle force production, Physiol Rev 88(4) (2008) 1243–76.

[4] D. Pye, J. Palomero, T. Kabayo, M.J. Jackson, Real-time measurement of nitric oxide in single mature mouse skeletal muscle fibres during contractions, J Physiol 581(Pt 1) (2007) 309–18.

[5] I.K. Aggeli, C. Gaitanaki, I. Beis, Involvement of JNKs and p38-MAPK/MSK1 pathways in H2O2-induced upregulation of heme oxygenase-1 mRNA in H9c2 cells, Cellular Signalling 18(10) (2006) 1801–12.

[6] S.G. Ahn, D.J. Thiele, Redox regulation of mammalian heat shock factor 1 is essential for Hsp gene activation and protection from stress, Genes & Development 17(4) (2003) 516–28.

[7] N. Bakkar, J. Wang, K.J. Ladner, H. Wang, J.M. Dahlman, M. Carathers, S. Acharyya, M.A. Rudnicki, A.D. Hollenbach, D.C. Guttridge, IKK/NF-kappaB regulates skeletal myogenesis via a signaling switch to inhibit differentiation and promote mitochondrial biogenesis, J Cell Biol 180(4) (2008) 787–802.

[8] M. Bar-Shai, E. Carmeli, A.Z. Reznick, The role of NF-kappaB in protein breakdown in immobilization, aging, and exercise: from basic processes to promotion of health, Ann N Y Acad Sci 1057 (2005) 431–47.

[9] G. Gloire, J. Piette, Redox regulation of nuclear post-translational modifications during NF-kappaB activation, Antioxid Redox Signal 11(9) (2009) 2209–22.

[10] H.S. Marinho, C. Real, L. Cyrne, H. Soares, F. Antunes, Hydrogen peroxide sensing, signaling and regulation of transcription factors, Redox Biology 2 (2014) 535–62.

[11] J. Zhang, G. Johnston, B. Stebler, E.T. Keller, Hydrogen peroxide activates NFkappaB and the interleukin-6 promoter through NFkappaB-inducing kinase, Antioxid Redox Signal 3(3) (2001) 493–504.

[12] M.C. Gomez-Cabrera, A. Salvador-Pascual, H. Cabo, B. Ferrando, J. Viña, Redox modulation of mitochondriogenesis in exercise. Does antioxidant supplementation blunt the benefits of exercise training?, Free Radic Biol Med 86 (2015) 37–46.

[13] H. Sies, Hydrogen peroxide as a central redox signaling molecule in physiological oxidative stress: Oxidative eustress, Redox Biology 11 (2017) 613–619.

[14] M.J. Jackson, A. McArdle, Age-related changes in skeletal muscle reactive oxygen species generation and adaptive responses to reactive oxygen species, J Physiol 589(Pt 9) (2011) 2139–45.

[15] H. Sies, Role of metabolic H2O2 generation: redox signaling and oxidative stress, J Biol Chem 289(13) (2014) 8735–41.

[16] B.K. Huang, H.D. Sikes, Quantifying intracellular hydrogen peroxide perturbations in terms of concentration, Redox Biology 2 (2014) 955–62.

[17] S. Stöcker, K. Van Laer, A. Mijuskovic, T.P. Dick, The Conundrum of Hydrogen Peroxide Signaling and the Emerging Role of Peroxiredoxins as Redox Relay Hubs, Antioxid Redox Signal 28(7) (2018) 558–573.

[18] Z.A. Wood, E. Schroder, J. Robin Harris, L.B. Poole, Structure, mechanism and regulation of peroxiredoxins, Trends in Biochemical Sciences 28(1) (2003) 32–40.

[19] B. Manta, M. Hugo, C. Ortiz, G. Ferrer-Sueta, M. Trujillo, A. Denicola, The peroxidase and peroxynitrite reductase activity of human erythrocyte peroxiredoxin 2, Archives of Biochemistry and Biophysics 484(2) (2009) 146–54.

[20] D. Talwar, J. Messens, T.P. Dick, A role for annexin A2 in scaffolding the peroxiredoxin 2-STAT3 redox relay complex, Nat Commun 11(1) (2020) 4512.

[21] R.M. Jarvis, S.M. Hughes, E.C. Ledgerwood, Peroxiredoxin 1 functions as a signal peroxidase to receive, transduce, and transmit peroxide signals in mammalian cells, Free Radic Biol Med 53(7) (2012) 1522–30.

[22] M.C. Sobotta, W. Liou, S. Stocker, D. Talwar, M. Oehler, T. Ruppert, A.N. Scharf, T.P. Dick, Peroxiredoxin-2 and STAT3 form a redox relay for H2O2 signaling, Nat Chem Biol 11(1) (2015) 64–70.

[23] T.J. Tavender, J.J. Springate, N.J. Bulleid, Recycling of peroxiredoxin IV provides a novel pathway for disulphide formation in the endoplasmic reticulum, EMBO Journal 29(24) (2010) 4185–4197.

[24] L. van Dam, M. Pagès-Gallego, P.E. Polderman, R.M. van Es, B.M.T. Burgering, H.R. Vos, T.B. Dansen, The Human 2-Cys Peroxiredoxins form Widespread, Cysteine-Dependent- and Isoform-Specific Protein-Protein Interactions, Antioxidants (Basel) 10(4) (2021).

[25] C. Henriquez-Olguin, R. Meneses-Valdes, P. Kritsiligkou, E. Fuentes-Lemus, From workout to molecular switches: How does skeletal muscle produce, sense, and transduce subcellular redox signals?, Free Radical Biology and Medicine 209 (2023) 355–365.

[26] C. Stretton, J.N. Pugh, B. McDonagh, A. McArdle, G.L. Close, M.J. Jackson, 2-Cys peroxiredoxin oxidation in response to hydrogen peroxide and contractile activity in skeletal muscle: A novel insight into exercise-induced redox signalling?, Free Radic Biol Med 160 (2020) 199–207.

[27] Q. Xia, J.C. Casas-Martinez, E. Zarzuela, J. Muñoz, A. Miranda-Vizuete, K. Goljanek-Whysall, B. McDonagh, Peroxiredoxin 2 is required for the redox mediated adaptation to exercise, Redox Biology 60 (2023) 102631.

[28] Q. Xia, P. Li, J.C. Casas-Martinez, A. Miranda-Vizuete, E. McDermott, P. Dockery, K. Goljanek-Whysall, B. McDonagh, Peroxiredoxin 2 regulates DAF-16/FOXO mediated mitochondrial remodelling in response to exercise that is disrupted in ageing, Mol Metab 88 (2024) 102003.

[29] M.J. Jackson, On the mechanisms underlying attenuated redox responses to exercise in older individuals: A hypothesis, Free Radic Biol Med 161 (2020) 326–338.

[30] K. Mamchaoui, C. Trollet, A. Bigot, E. Negroni, S. Chaouch, A. Wolff, P.K. Kandalla, S. Marie, J. Di Santo, J.L. St Guily, F. Muntoni, J. Kim, S. Philippi, S. Spuler, N. Levy, S.C. Blumen, T. Voit, W.E. Wright, A. Aamiri, G. Butler-Browne, V. Mouly, Immortalized pathological human myoblasts: towards a universal tool for the study of neuromuscular disorders, Skelet Muscle 1 (2011) 34.

[31] A. Bigot, A.F. Klein, E. Gasnier, V. Jacquemin, P. Ravassard, G. Butler-Browne, V. Mouly, D. Furling, Large CTG repeats trigger p16-dependent premature senescence in myotonic dystrophy type 1 muscle precursor cells, The American journal of pathology 174(4) (2009) 1435–42.

[32] M.M. Bradford, A rapid and sensitive method for the quantitation of microgram quantities of protein utilizing the principle of protein-dye binding, Analytical biochemistry 72(1-2) (1976) 248–254.

[33] C. Staunton, E. Owen, K. Hemmings, A. Vasilaki, A. McArdle, R. Barrett-Jolley, M. Jackson, Skeletal muscle transcriptomics identifies common pathways in nerve crush injury and ageing, Skeletal Muscle 12(1) (2022) 3.

[34] R. Kumar-Singh, J.S. Chamberlain, Encapsidated adenovirus minichromosomes allow delivery and expression of a 14 kb dystrophin cDNA to muscle cells, Human Molecular Genetics 5(7) (1996) 913–921.

[35] C. Staunton, E. Owen, N. Pollock, A. Vasilaki, R. Barrett-Jolley, A. McArdle, M. Jackson, HyPer2 imaging reveals temporal and heterogeneous hydrogen peroxide changes in denervated and aged skeletal muscle fibers in vivo, Scientific Reports 9(1) (2019) 14461.

[36] A. McArdle, D. Pattwell, A. Vasilaki, R. Griffiths, M. Jackson, Contractile activity-induced oxidative stress: cellular origin and adaptive responses, American Journal of Physiology-Cell Physiology 280(3) (2001) C621–C627.

[37] M. Martin, Cutadapt removes adapter sequences from high-throughput sequencing reads, EMBnet. journal 17(1) (2011) 10–12.

[38] B. Langmead, S.L. Salzberg, Fast gapped-read alignment with Bowtie 2, Nature methods 9(4) (2012) 357–359.

[39] M.I. Love, S. Anders, V. Kim, W. Huber, RNA-Seq workflow: gene-level exploratory analysis and differential expression, F1000Research 4 (2015).

[40] M. Flück, Functional, structural and molecular plasticity of mammalian skeletal muscle in response to exercise stimuli, J Exp Biol 209(Pt 12) (2006) 2239–48.

[41] K.S. Romanello, K.K. Teixeira, J.P.M. Silva, S.T. Nagamatsu, M.A.C. Bezerra, I.F. Domingos, D.A. Martins, A.S. Araujo, C. Lanaro, C.A. Breyer, Global analysis of erythroid cells redox status reveals the involvement of Prdx1 and Prdx2 in the severity of beta thalassemia, PLoS One 13(12) (2018) e0208316.

[42] J. Dalla Rizza, L.M. Randall, J. Santos, G. Ferrer-Sueta, A. Denicola, Differential parameters between cytosolic 2-Cys peroxiredoxins, PRDX1 and PRDX2, Protein Science 28(1) (2019) 191–201.

[43] X. Du, D.A. Williams, Interleukin-11: Review of Molecular, Cell Biology, and Clinical Use, Blood 89(11) (1997) 3897–3908.

[44] K. Zhao, Y. Yi, Z. Ma, W. Zhang, INHBA is a prognostic biomarker and correlated with immune cell infiltration in cervical cancer, Frontiers in Genetics 12 (2022) 705512.

[45] H. Yu, S. Zaveri, M. Pillai, H. Taluru, M. Schaible, S. Chaddha, A. Ahmed, S. Tfaili, P. Geraghty, The Role of Leukemia Inhibitory Factor in Counteracting the Immunopathology of Acute and Chronic Lung Inflammatory Diseases, Journal of Respiration 3(2) (2023) 86–100.

[46] S. Docherty, R. Harley, J.J. McAuley, L.A.N. Crowe, C. Pedret, P.D. Kirwan, S. Siebert, N.L. Millar, The effect of exercise on cytokines: implications for musculoskeletal health: a narrative review, BMC Sports Science, Medicine and Rehabilitation 14(1) (2022) 5.

[47] K. Vargas-Ortiz, V. Pérez-Vázquez, M.H. Macías-Cervantes, Exercise and sirtuins: a way to mitochondrial health in skeletal muscle, International journal of molecular sciences 20(11) (2019) 2717.

[48] J.P. Morton, A.C. Kayani, A. McArdle, B. Drust, The Exercise-Induced Stress Response of Skeletal Muscle, with Specific Emphasis on Humans, Sports Medicine 39(8) (2009) 643–662.

[49] L.B. Ivashkiv, L.T. Donlin, Regulation of type I interferon responses, Nature Reviews Immunology 14(1) (2014) 36–49.

[50] M.J. Jackson, C. Stretton, A. McArdle, Hydrogen peroxide as a signal for skeletal muscle adaptations to exercise: What do concentrations tell us about potential mechanisms?, Redox Biology (2020) 101484.

[51] O. Lyublinskaya, F. Antunes, Measuring intracellular concentration of hydrogen peroxide with the use of genetically encoded H(2)O(2) biosensor HyPer, Redox Biology 24 (2019) 101200.

[52] A. Philips, T. Cooper*, RNA processing and human disease, Cellular and Molecular Life Sciences CMLS 57 (2000) 235–249.

[53] A.V. Peskin, C.C. Winterbourn, The Enigma of 2-Cys Peroxiredoxins: What Are Their Roles?, Biochemistry (Mosc) 86(1) (2021) 84–91.

[54] S.G. Rhee, H.A. Woo, Multiple functions of peroxiredoxins: peroxidases, sensors and regulators of the intracellular messenger H₂O₂, and protein chaperones, Antioxid Redox Signal 15(3) (2011) 781–94.

[55] N.A. Strobel, J.M. Peake, A. Matsumoto, S.A. Marsh, J.S. Coombes, G.D. Wadley, Antioxidant supplementation reduces skeletal muscle mitochondrial biogenesis, Med Sci Sports Exerc 43(6) (2011) 1017–24.

[56] W. Szaflarski, M. Leśniczak-Staszak, M. Sowiński, S. Ojha, A. Aulas, D. Dave, S. Malla, P. Anderson, P. Ivanov, S.M. Lyons, Early rRNA processing is a stress-dependent regulatory event whose inhibition maintains nucleolar integrity, Nucleic Acids Res 50(2) (2022) 1033–1051.

[57] D. Shedlovskiy, J.A. Zinskie, E. Gardner, D.G. Pestov, N. Shcherbik, Endonucleolytic cleavage in the expansion segment 7 of 25S rRNA is an early marker of low-level oxidative stress in yeast, J Biol Chem 292(45) (2017) 18469–18485.

